# Antiviral innate immune memory in alveolar macrophages following SARS-CoV-2 infection

**DOI:** 10.1101/2023.11.24.568354

**Authors:** Alexander Lercher, Jin-Gyu Cheong, Chenyang Jiang, Hans-Heinrich Hoffmann, Alison W. Ashbrook, Yue S. Yin, Corrine Quirk, Emma J. DeGrace, Luis Chiriboga, Brad R. Rosenberg, Steven Z. Josefowicz, Charles M. Rice

## Abstract

Pathogen encounter results in long-lasting epigenetic imprinting that shapes diseases caused by heterologous pathogens. The breadth of this innate immune memory is of particular interest in the context of respiratory pathogens with increased pandemic potential and wide-ranging impact on global health. Here, we investigated epigenetic imprinting across cell lineages in a disease relevant murine model of SARS-CoV-2 recovery. Past SARS-CoV-2 infection resulted in increased chromatin accessibility of type I interferon (IFN-I) related transcription factors in airway-resident macrophages. Mechanistically, establishment of this innate immune memory required viral pattern recognition and canonical IFN-I signaling and augmented secondary antiviral responses. Past SARS-CoV-2 infection ameliorated disease caused by the heterologous respiratory pathogen influenza A virus. Insights into innate immune memory and how it affects subsequent infections with heterologous pathogens to influence disease pathology could facilitate the development of broadly effective therapeutic strategies.

## Introduction

Immune memory is critical to fend off and ameliorate pathology of recurring diseases caused by pathogens. This is not only beneficial for the individual, but also forms the basis of herd immunity and population health ^1^. Adaptive immune cells evolved to mount robust antigen-dependent responses and form long-lived memory cells. Complementary to this pathogen-specific immune memory, innate immune responses can facilitate the establishment of antigen-independent inflammatory memory ^2^. This innate immune memory is defined as an epigenetic memory-state of a cell following encounter with inflammatory cues that alters subsequent immune responses ^3,4^. The longevity of innate immune memory is influenced by cell type and stimulus and can be extended when established in immune progenitor cells in the bone marrow, tissue-resident stem cells or self-renewing tissue-resident macrophages ^5–8^. The live-attenuated tuberculosis vaccine Bacillus Calmette-Guérin (BCG) elicits innate immune memory in humans that lasts for months and reduces child mortality caused by heterologous infectious agents ^9–12^. Hence, innate immune memory is versatile and can provide long-lived cross-protection against heterologous pathogens. Still, how this phenomenon influences real-word infectious encounters, such as serial infections with heterologous pathogens, is poorly understood.

Respiratory pathogens have increased pandemic potential due to efficient airborne transmission and often widespread illness within populations. The recent pandemic caused by severe acute respiratory syndrome coronavirus 2 (SARS-CoV-2) and the resulting coronavirus disease 2019 (COVID-19) exemplified the far-reaching health and economic consequences of a new respiratory pathogen ^13^ encountering a highly permissive, immune naïve host population. Development of antigen-specific effective vaccines can help to achieve herd immunity requires time and success is not guaranteed. Additionally, viruses can rapidly evolve to escape antigen-specific immune memory ^14^. Intervention strategies built upon the antigen-independent nature of innate immune memory could provide increased robustness to achieve protective immunity in a naïve population ^4^. Alveolar macrophages are the most abundant immune cell type in the airway and form a stem-like immune cell population that is the first line of defense against respiratory pathogens ^15–17^. Following bacterial or viral infections, alveolar macrophages are capable of forming innate immune memory that can affect the outcome of secondary lung diseases, such as bacterial pneumonia or cancer ^8,18,19^. There is a gap in our understanding of innate immune memory in the context of commonly circulating respiratory viruses. Specifically, how past infections impact subsequent infections with unrelated viruses via epigenetic imprinting across airway-resident immune cell lineages remains understudied. Such insights into the establishment, maintenance, and recall of innate immune memory may therefore facilitate the development of novel therapeutic strategies that target a broad range of respiratory pathogens. In this study, we used a disease-relevant murine infection model of SARS-CoV-2 recovery to study innate immune memory in airway-resident immune cells at single cell resolution and determine how this influences infection outcome of secondary influenza A virus infection.

## Results

### Past SARS-CoV-2 infection leads to establishment of epigenetic imprinting in alveolar macrophages

We intranasally (i.n.) infected C57Bl/6J wild type mice with 6,000 PFU of the mouse-adapted strain MA10 of SARS-CoV-2 (SARS2) ^20^. Upon SARS2 infection, mice transiently lose body weight loss, which they regain by 20-30 days post infection (dpi) (Figure 1A). SARS2 is cleared by 15 dpi ^20,21^ and we confirmed absence of SARS2 RNA and antigen in lung tissue via RT-qPCR or histology at 30 dpi (Figures S1A and S1B). Gene expression levels of many antiviral (*Ifit1*, *Bst2*, *Ifitm3*, *Isg15*) and inflammatory genes (*Tnfa*, *Il6*) were not significantly different in bulk lung tissue of recovered vs. naïve animals (Figure S1C). Immune cell profiling of bronchoalveolar lavage fluid (BALF) via flow cytometry revealed no significant differences in numbers of alveolar macrophages (CD45^+^CD11c^+^SiglecF^+^), NK cells (CD45^+^CD11c^−^SiglecF^−^NK1.1^+^) or neutrophils (CD45^+^CD11c^−^ SiglecF^−^CD11b^+^Ly6G^+^) in recovered vs. naïve mice (Figure S1D). The inflammatory milieu in recovered and naïve airways was comparable. Among 32 detectable cytokines, only minor differences were observed for IFN-γ, IL-2, IL-13 (decreased) and CXCL9, CCL19, CCL22, TIMP-1 (increased) in recovered vs. naïve BALF (Figures S1E and S1F). Flow cytometric analyses of major subsets of bone marrow progenitor cells were comparable between recovered and naïve mice (Figure S1G). CD4 and CD8 T cell numbers were significantly increased in recovered BALF and expressed surface markers associated with effector memory (CD44^+^CD62L^−^) or tissue-resident memory (CD69+CD103^+^) cells (Figure S1H). Together, these data show that SARS2-recovered animals do not retain active inflammatory responses or tissue pathology.

**Figure 1:**
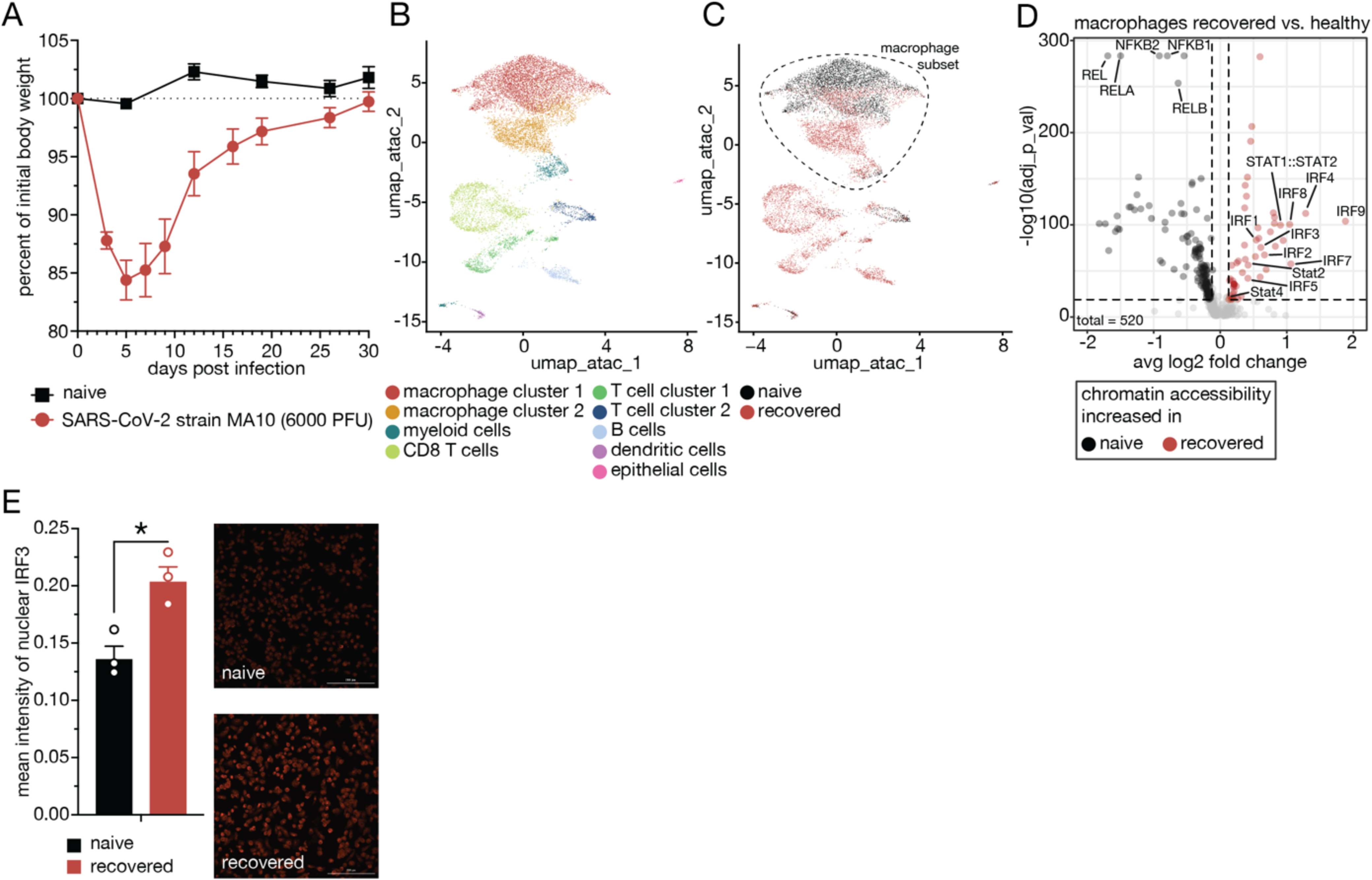
Past SARS-CoV-2 infection establishes epigenetic memory in airway-resident macrophages. (A) Body weights of naive and SARS-CoV-2 (strain MAI 0) infected C57BI/6J mice, n = 11 -12. (B) UMAP clustering of single nuclei combined ATAC/RNA-seq data (10x Multiome) and annotated cell clusters of airway-resident cells from naive and SARS-CoV-2-recovered mice based on ATAC-seq data, n = 3. (C) Recovered and naive sample annotation of UMAP cluster­ing (B) with dashed line indicating the macrophage subset. (D) TF motif-associated chromatin accessibility analyses of recov­ered vs. naive sub-setted macrophages. (E) Quantification of mean fluorescent intensity of nuclear IRF3 in airway-resident macrophages isolated from naive and SARS-CoV-2-recovered animals and representative image, n = 3. Data are mean ± s.e.m. n values indicate the number of mice or replicates. For (D), statistical analysis was performed using Wilcoxon’s test. For (E), statistical analysis was performed using Student’s t-test with Bonferroni correction when multiple comparisons were performed. *p < 0.05; **p < 0.01; ***p < 0.001.

Next, we tested whether past SARS2 infection leads to sustained cell-intrinsic changes of airway-resident cells. We investigated changes of chromatin accessibility and transcript levels of individual cells in recovered (31 dpi) or naïve BALF (n = 3) via single nuclei combined ATAC/RNA sequencing (10x Chromium Single Cell Multiome ATAC + Gene Expression). We profiled 13,622 single nuclei (5,669 naïve and 7,339 recovered) and identified major clusters based on chromatin accessibility using Seurat ^22^ (Table S1). Most profiled nuclei were derived from macrophages (65%) that clustered as two separate populations, followed by CD8 T cells (16%) (Figure 1B). In accordance with flow cytometry data, CD8 T cells were primarily present in BALF of recovered mice (Figure 1C). Intriguingly, macrophage clusters 1 and 2 were mainly driven by experimental condition, with macrophage cluster 2 consisting almost exclusively of nuclei isolated from recovered animals (Figure 1C). To compare epigenetic changes in recovered and naïve macrophages following SARS2 infection, we extracted and re-clustered all putative macrophages from the dataset, including myeloid cells (Figures 1S and S1I). Differentially regulated genes (DEG) were enriched for gene ontology (GO) terms related to cytokine production and myeloid cell differentiation for naïve and histone modification and chromatin organization in recovered macrophages (Figure S1J). Yet, the transcriptomic profiles of recovered and naïve macrophages were surprisingly comparable with only 36 of 5,701 detected genes being significantly regulated by more than 1.4-fold between conditions (Figure S1K; Table S1). However, comparing chromatin accessibility transcription factor (TF) binding motifs using chromVAR ^23^ revealed distinct differences in recovered and naïve macrophages. Recovered macrophages showed increased accessibility of TF binding motifs associated with antiviral immune response and type I interferon (IFN-I) signaling, including interferon regulatory factors (IRFs) and signal transducer and activator of transcription (STAT) proteins (Figure 1D). In contrast, accessibility of binding motifs associated with nuclear factor kappa-light-chain-enhancer of activated B cell (NF-κB) was lower in recovered macrophages (Figure 1D; Table S1). Immunofluorescence staining of airway-resident macrophages *ex vivo* showed that recovered cells had increased nuclear levels of IRF3 but not RELA (p65 subunit of NF-κB) relative to naïve cells (Figures 1E and S1L).

These data show that alveolar macrophages in mice retain epigenetic imprinting following SARS2 infection.

### Past COVID-19 leads to establishment of epigenetic imprinting in circulating monocytes in patients

To determine if an antiviral program persists in humans following SARS2 infection and clearance, we recruited a patient cohort during the initial infection wave in 2020, prior to availability of COVID-19 vaccines. We performed single nuclei combined ATAC/RNA sequencing on peripheral blood mononuclear cells (PBMCs) isolated from patients recovered from mild COVID-19 after 2-4 months (n = 3) and healthy controls (n = 7). Based on transcriptional profile, we identified major cell clusters in PBMCs (Figure 2A). We focused our analyses on myeloid cell clusters (CD14^+^ and CD16^+^ monocytes and dendritic cells (DCs)) and found increased expression of genes related to the GO category “Defense Response to Virus” in recovered patients (Figures 2B and S2A, Table S2). Although to a lesser extent, this gene module was also enriched in recovered murine macrophages in BALF (Figure 2C). Gene module enrichment analyses of DEG in murine recovered vs. naïve BALF macrophages (Figure S1I-S1K) underscored human CD14^+^ monocytes as suitable cell population to analyze epigenetic memory in patients (Figures 2D and S2B). Consistent with data obtained from murine macrophages in BALF, TF binding site accessibility was significantly higher for IRFs and lower for NF-κB in CD14^+^ monocytes from recovered compared to healthy patients (Figure 2E and Table S2).

**Figure 2:**
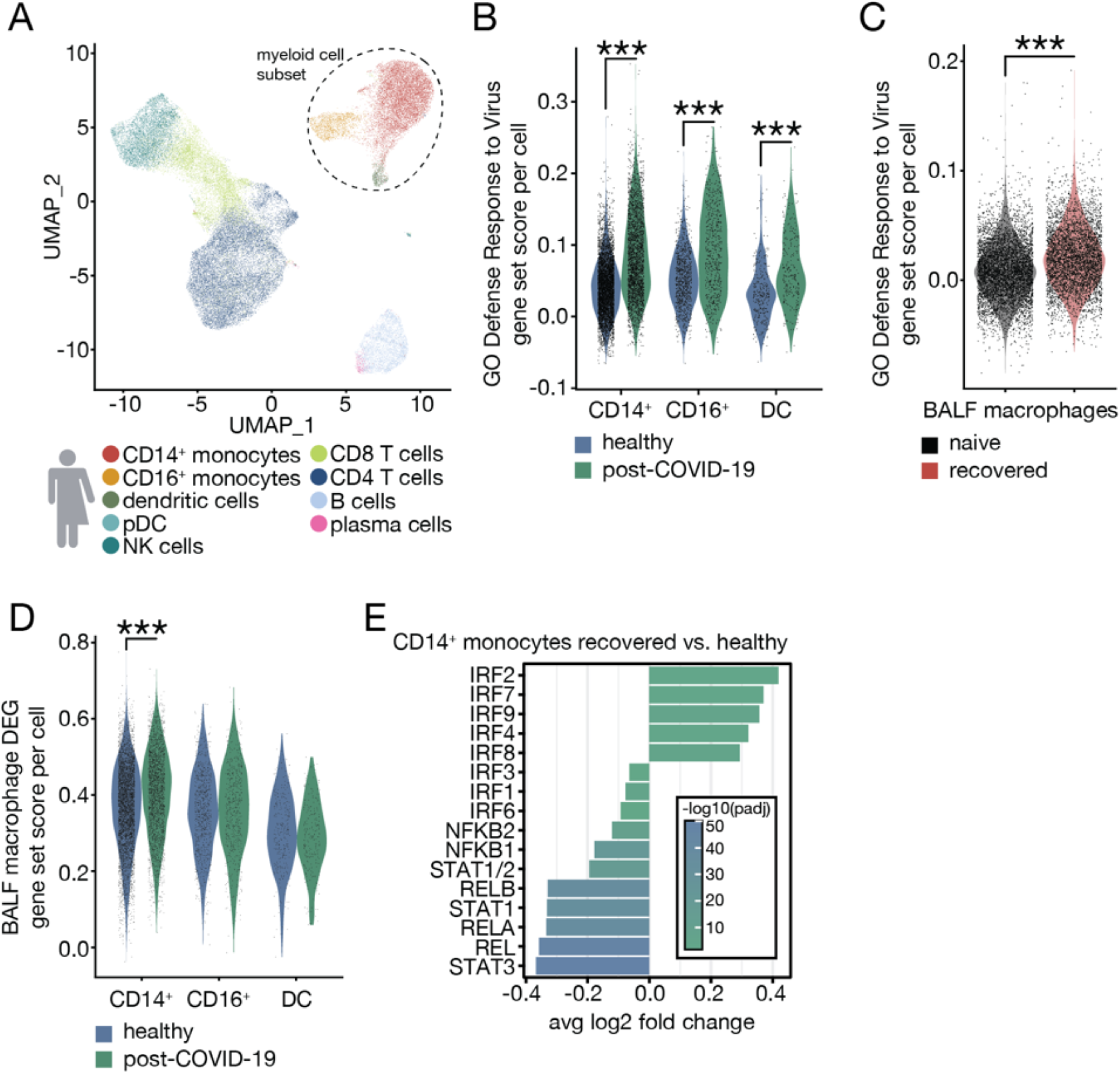
Past COVID-19 establishes epigenetic memory in circulating monocytes in patients. (A) UMAP clustering of single nuclei combined ATAC/RNA-seq data and annotated cell clusters of peripheral blood mononu­clear cells (PBMCs) of recovered (2-4 months) mild COVID-19 and healthy patients with dashed line indicating the myeloid cell subset, n = 3-7. (B-C) Gene set expression score of GO: Defense Response to Virus in recovered and healthy circulating human myeloid cells (CD14+ and GD16+ monocytes and dendritic cells) (B) and murine BALE macrophages (C). (D) Gene set expression score of differentially expressed gene (DEG) module in circulating human myeloid cells. (E) Significantly different accessible TF motif-associated chromatin of recovered vs. healthy CD14+ monocytes. Data are represented as violin plots with each dot corresponding to one individual cell. For (B-E), statistical analyses were performed using Wilcoxon’s test, “p < 0.05; *‘p< 0.01; *“p < 0.001.

### Past SARS2 infection increases secondary antiviral immune responses in murine airway-resident macrophages

To investigate whether SARS2-associated epigenetic imprinting alters secondary immune responses, we isolated airway-resident macrophages from recovered and naïve animals and stimulated them *ex vivo* with the synthetic viral double-stranded RNA mimic polyinosinic-polycytidylic acid (polyIC). Upon polyIC stimulation, recovered macrophages displayed significantly higher levels of nuclear IRF3 than naïve macrophages (Figure 3A). At the transcriptional level, control and polyIC-stimulated recovered and naïve macrophages clustered by stimulation and infection history (Figure S3A). We identified 2,654 DEG and hierarchical clustering revealed both infection history-specific gene sets (clusters 1 and 6) and polyIC response genes, including interferon-stimulated genes (ISGs) ^24^ (cluster 2) (Figure 3B; Table S3). Genes associated with naïve cells were enriched for GO terms related to lipid and cholesterol metabolism (Figure 3C), whereas recovered cells expressed genes related to macrophage activation and inflammatory response (Figure 3D). Cluster 2 was enriched for GO terms related to antiviral immunity and interferon response (Figure 3E). Recovered macrophages exhibited a pronounced hyperresponsiveness to polyIC, characterized by robust induction of ISGs, including *Ifit1*, *Ifitm3*, *Bst2* (Figures S3B-S3D; Table S3). As a functional consequence, recovered macrophages were significantly less susceptible to infection with vesicular stomatitis virus expressing a GFP reporter (VSV-GFP) compared to naïve macrophages (Figure 3F). These findings suggest that SARS2-associated epigenetic imprinting of airway-resident macrophages results in innate immune memory that augments subsequent antiviral immune responses.

**Figure 3:**
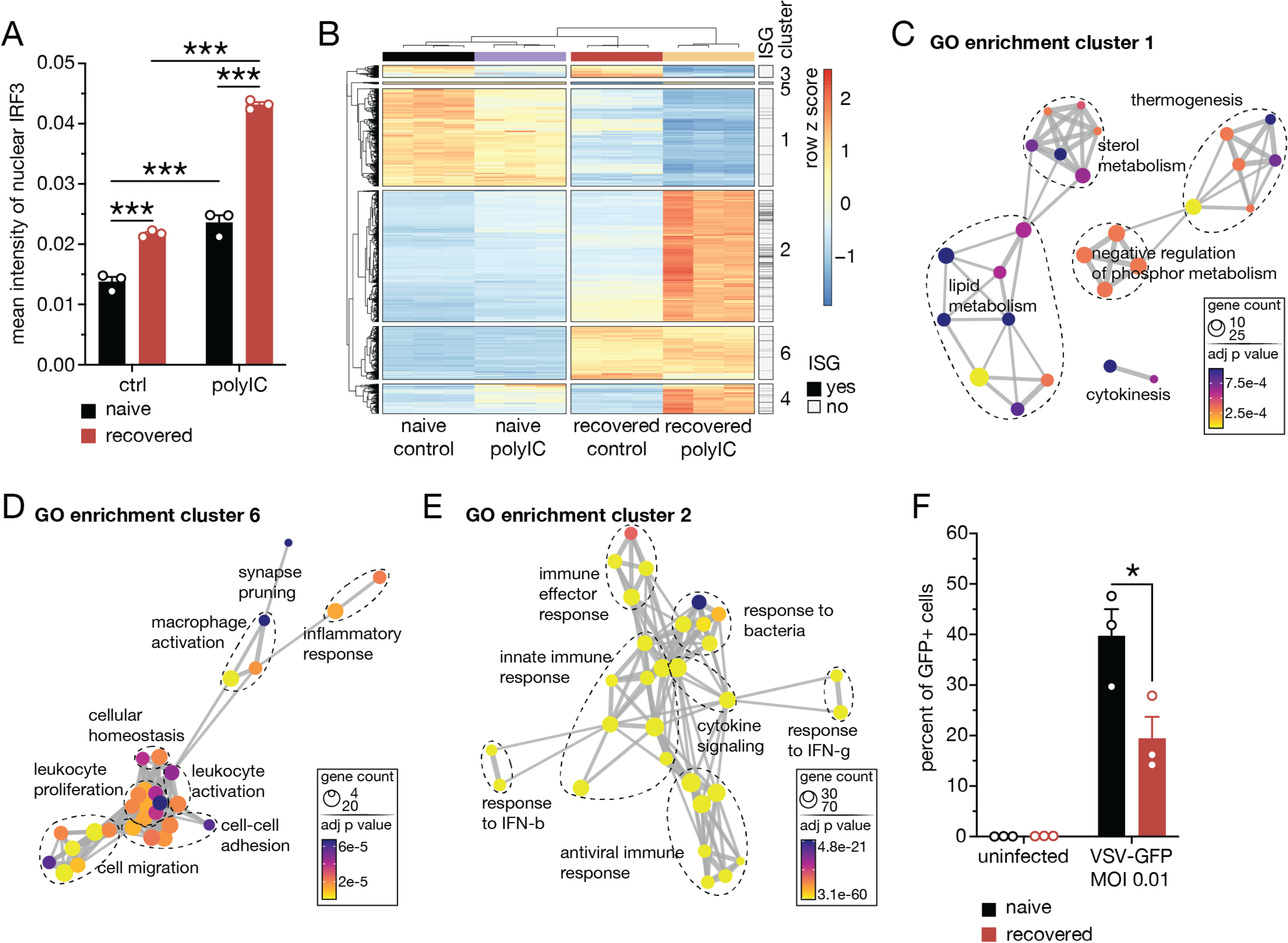
Past SARS-CoV-2 infection leads to increased secondary antiviral responses in airway-resident macro­phages. (A) Quantification of mean fluorescent intensity (MFI) of IRF3 in control- or polylC-stimulated airway-resident macrophages isolated from naive and SARS-CoV-2-recovered animals after 24 hours, n = 3. (B) Hierarchical clustering of differentially expressed genes (DEG) of control- or polylC-stimulated airway-resident macrophages isolated from naive or SARS-CoV-2-re-covered animals after 6 hours, n = 3. (0-E), Gene ontology (GO) enrichment analyses of genes in clusters 1 (0), 6 (D), 2 (E). (F) Percent of VSV-GFP infected airway-resident macrophages isolated from naive or SARS-CoV-2-recovered animals, n = 3. For (A and F), statistics were calculated using Student’s t-test with Bonferroni correction when multiple comparisons were performed. For (0-E), hypergeometric p values were adjusted for multiple testing with Benjamini-Hochberg correction. *p < 0.05; **p < 0.01; ***p < 0.001.

### Viral PAMP encounter is sufficient to establish antiviral innate immune memory in primary alveolar macrophages and requires canonical IFN-I signaling

To explore these findings using a tractable *in vitro* system, we employed a long-term culture system of primary alveolar macrophages ^25^. We exposed alveolar macrophages to polyIC for 24h and then re-stimulated them with polyIC after 5 days. Alveolar macrophages that previously experienced polyIC showed significantly increased nuclear IRF3 levels upon restimulation with polyIC (polyIC/polyIC) compared to control-experienced cells (control/polyIC) (Figure 4A). Like SARS2-experienced alveolar macrophages, we did not observe significant differences in nuclear localization of RELA (p65) (Figure S4A). Differential transcriptomic analyses confirmed a significantly more robust antiviral recall response (Figures 4B and 4C; Table S4) and ISG induction (*Ifit1*, *Ifitm3* and *Bst2*) (Figures S4B-S4D) in polyIC/polyIC vs. control/polyIC cells. Functionally, this correlated with a 17-fold increased resistance to infection with VSV-GFP as assayed after 5 days and was maintained up to 14 days, albeit to a lesser extent (Figures S4E and S4F). Memory formation required IFN-I signaling during initial polyIC exposure (Figure 4D) and the canonical downstream transcription factor IRF9 (Figure 4E). Mice that received polyIC-experienced alveolar macrophages 3 days prior to infection showed significantly ameliorated body weight loss upon infection with influenza A/PR/8/34 virus (PR8) (Figure 4F). Thus, like SARS2 infection *in vivo*, polyIC exposure *in vitro* leads to innate immune memory formation in alveolar macrophages that is sufficient to ameliorate pathology of a viral infection *in vivo*.

**Figure 4:**
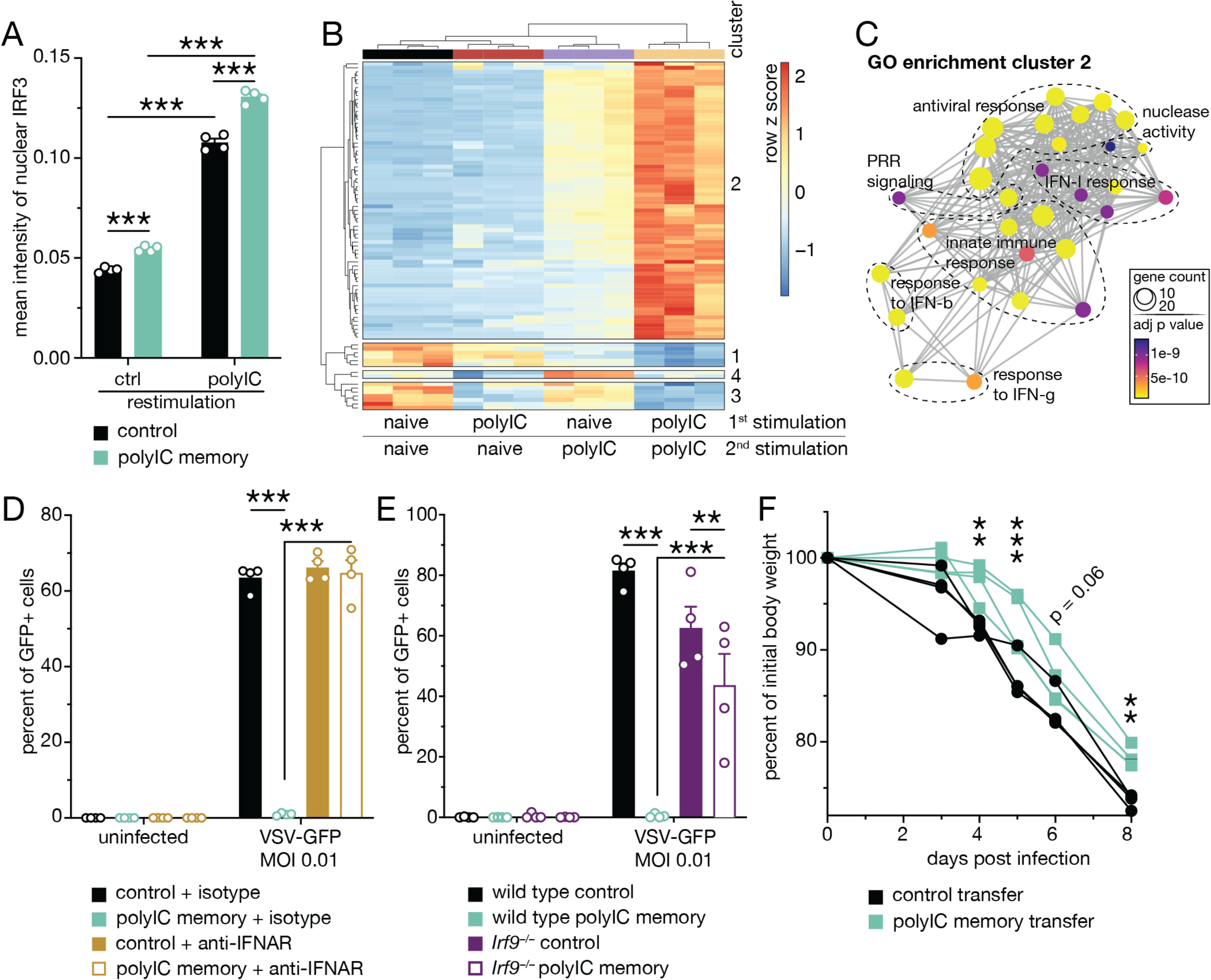
Viral PAMP exposure is sufficient to establish innate immune memory in alveolar macrophages in vitro. (A) Quantification of mean fluorescent intensity (MFI) of IRF3 in control- or polylC-stimulated control- or polylC-experienced in vitro cultured alveolar macrophages after 24 hours, n = 4. (B) Hierarchical clustering of differentially expressed genes (DEG) of polylC/polylO vs. control/polylO alveolar macrophages 6 hours after re-stimulation, n = 3. (0) Gene ontology (GO) enrichment analyses of genes in clusters 2. (D) Percent of VSV-GFP infected control- or polylC-experienced alveolar macrophages with or without anti-IFNAR blocking antibody treatment during initial polylC exposure, n = 4. (E) Percent of VSV-GFP infected control- or polylC-experienced wild type or IrfQ—/— alveolar macrophages, n = 4. (F) Body weights of influenza A/PR/8/34 virus infected C57BI/6J mice following transfer of control- or polylC-experienced alveolar macrophages, n = 4. Data are mean ±s.e.m. n values indicate the number of mice or replicates. For (A), statistics were calculated using Student’s t-test with Bonferroni correction when multiple comparisons were performed. For (C), hypergeometric p values were adjusted for multiple testing with Benjamini-Hochberg correction. For (D-E), statistical analysis was performed using Two-Way ANOVA comparison with Bonferroni correction. For (F), statistical analysis was performed using Two-Way ANOVA comparison with Bonferroni correc­tion. *p < 0.05; **p < 0.01; ***p < 0.001.

### Past SARS-CoV-2 infection can ameliorate the pathology of secondary influenza A virus infection

Finally, we investigated whether innate immune memory following SARS2 infection affects disease pathology caused by a heterologous respiratory virus. We challenged naïve or SARS2-recovered animals with a sub-lethal dose of PR8 (naïve/PR8 and SARS2/PR8, respectively) and found significantly reduced body weight loss in SARS2/PR8 animals (Figure S5A). Likewise, past SARS2 infection ameliorated body weight loss and reduced lethality using a higher (LD50) infectious dose of PR8 (Figures 5A and 5B). Immune cell profiling of BALF revealed significantly reduced neutrophil numbers in the SARS2/PR8 group but showed no significant differences in alveolar macrophages or NK cell numbers (Figures 5C-5E). Increased CD8 T cell numbers in SARS2-recovered animals (Figure S5H) were maintained in the early phase of PR8 infection (3dpi), but not at later time points (Figure 5F). Hyperinflammatory responses in the airway are the major determinant of lethal influenza infection ^26^. Several pro-inflammatory cytokines and chemokines (IL-1β, IL-17, CCL4 and CCL12) were significantly reduced in BALF of SARS2/PR8 compared to naïve/PR8 animals at 5 dpi (Figure 5G), suggesting that past SARS2 infection limits dysregulated inflammatory responses. Notably, there was no significant difference in viral RNA levels in lung tissue between naïve/PR8 and SARS2/PR8 animals (Figure S5B).

**Figure 5:**
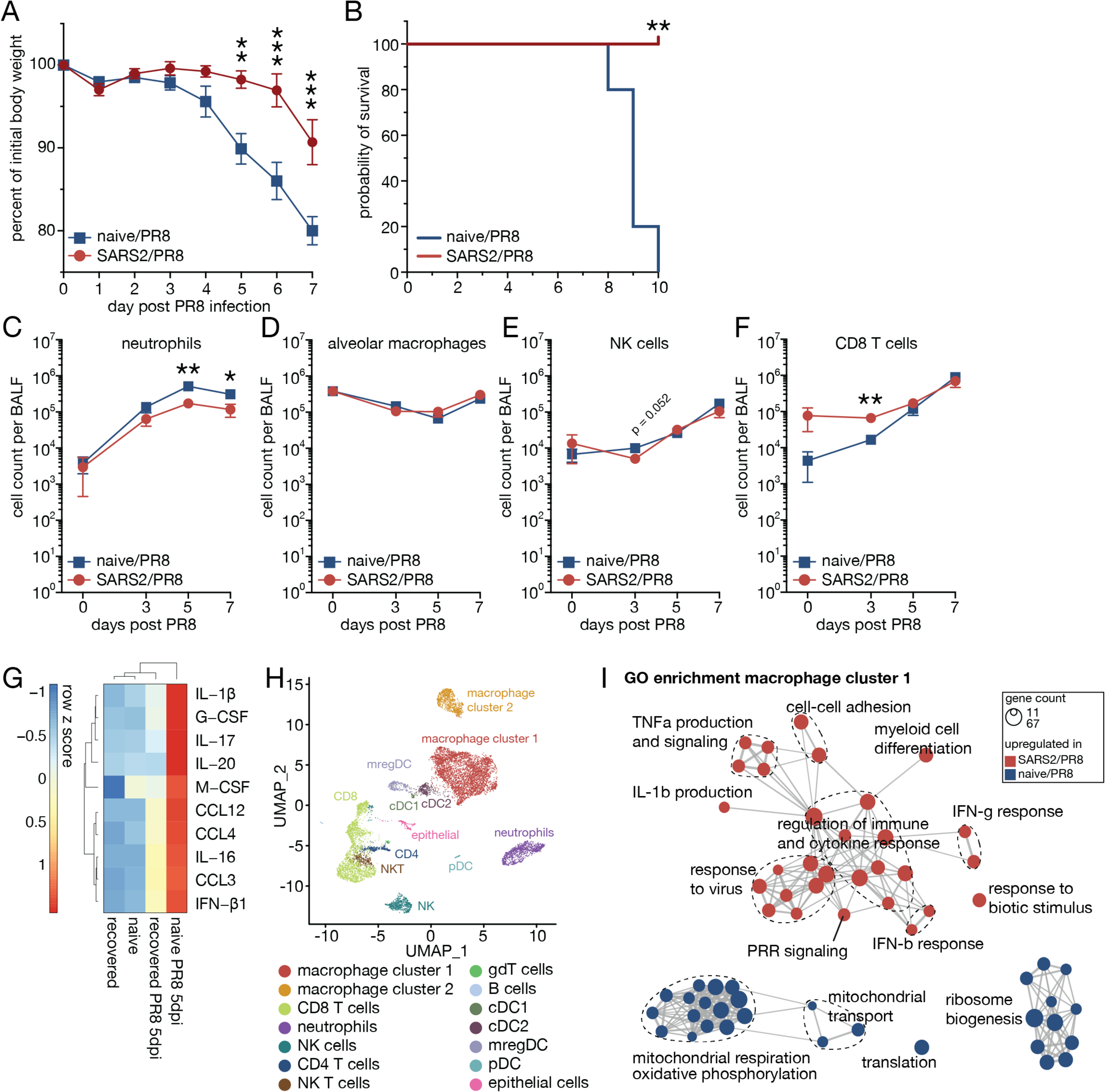
Past SARS-CoV-2 infection ameliorates secondary influenza A virus infection. (A) Body weights of naive and SARS-CoV-2-recovered animals infected with influenza A/PR/8/34 virus at LD50 (naïve/PR8 or SARS2/PR8). n = 5. (B) Survival percentages of naïve/PR8 and SARS2/PR8 animals, n = 5. (C-F) Kinetics of neutrophils (0), alveolar macrophages (D), NK cells (E) and 0D8 T cells (F) in bronchoalveolar lavage fluid (BALF) of naïve/PR8 and SARS2/PR8 animals, n = 3-7. (G) Significantly different cytokines and chemokines in BALF of naïve/PR8 and SARS2/PR8 animals at 5 days after PR8 infection, n = 4-5. (H) UMAP clustering and major cell cluster annotation of single cell RNA-seq data of BALF from naïve/PR8 and SARS2/PR8 animals at 7 days post PR8 infection. (I) Gene ontology (GO) enrichment analyses of genes associated with cells isolated from naïve/PR8 or SARS2/PR8 BALF in macrophages cluster 1 (G). Data are mean ±s.e.m. n values indicate the number of mice or replicates. For (A), statistical analysis was performed using Two-Way ANOVA comparison with Bonferroni correction. For (B), statistical analysis was performed using a log-rank Mantel-Cox test. For (C-G), statistics were calculated using Student’s t-test with Bonferroni correction when multiple comparisons were performed. For (I), hypergeometric p values were adjusted for multiple testing with Benjamini-Hochberg correction. *p < 0.05; **p < 0.01; ***p < 0.001.

To identify cell type-specific and cell-intrinsic differences associated with secondary infection, we performed single cell RNA-seq (scRNA-seq) of naïve/PR8 and SARS2/PR8 BALF at 7dpi. We sequenced 12,032 single cells (5,004 naïve/PR8 and 7,028 SARS2/PR8) and identified major cell clusters in which the majority were macrophages (40%), followed by CD8 T cells (18%) (Figure 5H). Macrophages separated into clusters 1 (82%) and 2 (18%) characterized by increased antiviral immunity and fatty acid metabolism, respectively (Figure S5C; Table S5). Cells in macrophage cluster 1 and neutrophils expressed particularly high levels of ISGs (Figure S5D). Within those clusters, DEG analyses revealed significantly increased expression of antiviral genes in cells isolated from SARS2/PR8 compared to naïve/PR8 animals (Figures 5I and S5E; Table S5). DEG in CD8 T cells were enriched for GO terms related to T cell differentiation and RNA metabolism (SARS2/PR8) and oxidative phosphorylation and ATP metabolism (naïve/PR8), suggesting no antigen-independent bystander activation of SARS2-specific cells in SARS2/PR8 animals (Figure S5F; Table S5). These data highlight a beneficial effect of past SARS2 infection on disease pathology caused by the heterologous pathogen influenza A PR8.

## Discussion

Antigen-independent, innate immune memory alters secondary inflammatory responses. Consequently, heterologous pathogens can indirectly influence each other, including emerging pathogens ^4^.

Innate immune memory can be established locally or centrally. Systemic exposure to PAMPs can result in epigenetic imprinting of immune progenitor cells in the bone marrow and central innate immune memory that facilitates tissue cross-protection ^6,7,27–29^. Further, innate immune memory mounted by long-lived stem cells can provide extended longevity compared to short-lived effector immune cells ^5,30^. There is evidence of durable epigenetic memory in hematopoietic stem and progenitor cells and circulating monocytes from human patients following severe SARS-CoV-2 infection and hospitalization that translates to altered secondary responses ^31^. Yet, pre-clinical animal models are critical to investigate the quality and breadth of SARS-CoV-2-induced innate immune memory – particularly at the tissue level and in the context of mild disease. The murine infection model of SARS-CoV-2 strain MA10 recapitulates many features of COVID-19 in humans ^20^. We discovered that past SARS-CoV-2 infection induces local innate immune memory via epigenetic imprinting of airway-resident macrophages. Alveolar macrophages form a self-renewing macrophage population of the airway that maintains itself throughout life at an estimated cell division rate of more than 2 weeks ^16,32^. Hence, epigenetic remodeling in airway-resident macrophages may contribute to the establishment of sustained, local innate immune memory. Of note, severe lung injury can coincide with depletion of alveolar macrophages that are sustainably replenished by infiltrating monocytes ^15,33^. These monocyte-derived alveolar macrophages can display an inflammatory phenotype and contribute to innate immune memory on a population level ^34,35^.

Following SARS-CoV-2, epigenetic memory of airway-resident macrophages was strongly skewed towards increased chromatin accessibility of IRF-related transcription factors. This allowed enhanced and rapid secondary antiviral responses, limiting detrimental hyperinflammatory responses in secondary influenza virus A infection. Epigenetic and transcriptional profiling of myeloid cells in a cohort comprised of healthy individuals and patients recovered from mild COVID-19 before availability of COVID-19 vaccine corroborated these findings. Consistent with our observations, circulating CD14^+^ monocytes from individuals who received a trivalent seasonal influenza vaccine exhibited increased chromatin accessibility for IRFs and antiviral gene expression ^36^.

Next to epigenetic imprinting, recovered macrophages retained an altered metabolic transcriptional program that suggests reduced activity of fatty acid metabolism. A similar metabolic switch is associated with increased inflammatory responses of immune cells ^37–39^ and is in agreement with increased glycolytic activity of alveolar macrophages in adenovirus-recovered animals ^8^. Interestingly, altered secondary immune responses of LPS-experienced alveolar macrophages seem to depend on fatty acid oxidation rather than glycolysis ^18^.

Similar to bacterial PAMP exposure ^18,40^, the synthetic viral dsRNA mimic polyIC was sufficient to establish innate immune memory in alveolar macrophages that depended on canonical IFN-I signaling and the downstream TF IRF9. This aligns with our finding of increased chromatin accessibility of the IRF9 binding motif in airway-resident macrophages following SARS-CoV-2 infection. Some studies suggest that IFN-I is sufficient to establish epigenetic memory, but appears to be more robust in immune cells compared to fibroblasts ^18,41^. Exposure of alveolar macrophages to polyIC led to increased secondary antiviral responses *in vitro* and ameliorated disease caused by influenza A virus *in vivo*. These findings show that viral PAMP recognition can lead to formation of innate immune memory in alveolar macrophages that is sufficient to influence pathology of secondary respiratory viral infections. Our description of matching epigenetic antiviral imprinting in myeloid cells of recovered COVID-19 patients implies that these shared programs might broadly modulate human responses to subsequent infections.

Further, innate immune memory in macrophages can impact on other immune cells and shape adaptive T cell responses ^8,42^. While we did not observe pronounced differences in T cell recruitment and transcriptional profile between SARS2/PR8 and naïve/PR8 animals, we found increased ISG expression in alveolar macrophages and neutrophils. These data suggest that alveolar macrophages in SARS2/PR8 animals retain elevated antiviral activity during the acute phase of PR8 infection. Increased expression of ISGs in recruited neutrophils may be a secondary effect resulting from disparate inflammatory conditions in the airways of naïve/PR8 and SARS2/PR8 animals.

Innate immune memory can be pro-inflammatory or tolerogenic which is thought to be primarily determined by initial PAMP encounter ^43^. Increased LPS levels dampen secondary inflammatory responses in monocytes ^43,44^. In line with this, monocytes of sepsis-recovered patients displayed immune paralysis and impaired phagocytic activity ^45^. While these tolerogenic recall responses are detrimental in certain secondary diseases, including cancer ^46^, they might also foster microbe-host co-existence and limit immunopathology ^47^. LPS-induced tolerance in monocytes can be reverted by subsequent exposure to the fungal PAMP β-glucan ^44^. This highlights a therapeutic opportunity, but also emphasizes that crosstalk between distinct inflammatory cues shapes the quality of innate immune memory in a cell-autonomous fashion. Likewise, the nature of secondary inflammatory signals influences the robustness of recall responses. Previous BCG encounter ameliorates disease caused by the respiratory pathogen influenza A virus but not SARS-CoV-2 and might be linked to differences in pulmonary vasculature damage and pathogen dissemination ^28^. These observations highlight innate immune memory as a complex dynamic trait. Especially in the context of infectious diseases, individual contributions and interplay of pathogen, host immune response, tissue damage and cellular heterogeneity remain incompletely understood ^8,18,19,33,35,40^. Well-controlled clinically relevant model systems are key to systematically dissect the interplay of inflammatory cues shaping the quality of innate immune memory.

Innate immune memory is not limited to infectious diseases but is also established in autoimmune disorders. Chronic rhinosinusitis in humans leads to epigenetic remodeling in basal stem cells of the upper airway that impacts inflammatory responses ^48^. Allergic asthma causes inflammatory imprinting in macrophages that exacerbates disease in a TNF-dependent fashion ^49^. Further, tissue crosstalk in autoimmune-induced innate immune memory can shape pathology of arthritis following recovery from periodontitis ^50^. There is a gap of knowledge regarding the crosstalk of pathogen and autoimmune-associated innate immune memory, as well as the reciprocal influence of disease pathology.

This study shows that past SARS-CoV-2 infection leads to establishment of epigenetic memory of airway-resident macrophages. Formation of innate immune memory depended on viral PAMP sensing and IFN-I signaling. This promoted increased secondary antiviral responses that translated to ameliorated pathology caused by subsequent challenge with influenza A virus. One can imagine that in the real-world onslaught of respiratory pathogens, the induction and duration of antigen-independent innate immune memory, along with the timing and nature of pathogen exposure, will have significant impact on infection outcome.

## Acknowledgements

The authors would like to thank Roni Winkler, Michael Bauer, Margaret MacDonald, Shira Weingarten-Gabbay, Tyler Lewy, Hsuan-An Chen, Mariana Nogueira Batista and the Rice lab for valuable input on experimental design, data interpretation and feedback on the manuscript. We thank Ralph Baric for sharing the mouse-adapted SARS-CoV-2 strain MA10 and the Flow Cytometry Resource Center at Rockefeller University for technical support.

## Data availability

RNA sequencing and 10x Multiome data will be made publicly available once the manuscript is published.

## Author contributions

A.L. and C.M.R. wrote the manuscript. A.L. designed and performed *in vivo* and *in vitro* studies. A.L., J.G.C., B.R.R. performed bioinformatic analyses. C.J., H.H.H., A.W.A., Y.S.Y., E.J.D. performed *in vitro* studies. C.Q. supported the *in vivo* studies. L.C. performed histological analyses. A.L., B.R.R., S.Z.J., C.M.R. designed and analyzed experiments. C.M.R. supervised the study. All co-authors provided feedback on the manuscript.

## Material and methods

### Mice

C57Bl/6J wild type mice were obtained from the Jackson Laboratory (strain #000664). *Irf9^−/–^* mice (RBRC00916) ^51^ were obtained from Riken Bioresource Center, Japan. Mice were maintained and bred at the AAALAC-accredited Comparative Bioscience Center of the Rockefeller University. All mouse experiments were in accordance with the NIH Guide for the Care and Use of Laboratory Animals and approved by the Institutional Animal Care and Use Committee of Rockefeller University. Four-to six-month-old mice of both sexes were used for SARS-CoV-2 strain MA10 infection experiments. For all other experiments, adult mice (older than 2 months) of both sexes were used. For survival studies, body weight loss greater than 20% of initial weight was defined the humane endpoint. All SARS-CoV-2 animal experiments and downstream processing of live cells non-inactivated tissues were conducted under biosafety level 3 (BSL-3) containment in compliance with institutional and federal guidelines.

### Cell lines

VeroE6 cells (*Chlorocebus sabaeus*; sex: female, kidney epithelial) obtained from the ATCC (CRL-1586) and Ralph Baric (University of North Carolina at Chapel Hill), and Huh-7.5 hepatoma cells (*Homo sapiens*; sex: male, liver epithelial) ^52^ were cultured in Dulbecco’s Modified Eagle Medium (DMEM, Thermo Fisher Scientific #11995065) supplemented to contain 1 % non-essential amino acids (NEAA, Thermo Fisher Scientific #11140076) and 10 % fetal bovine serum (FBS, HyClone Laboratories, Lot. #AUJ35777). BHK-21 cells (*Mesocricetus auratus*, sex: male, fibroblast) were obtained from ATCC (#CCL-10) and MDCK cells (*Canis familiaris*, sex: female, kidney epithelial) were obtained from ATCC (#CCL-34) and cultured in Modified Eagle Medium (MEM, Thermo Fisher Scientific #11095080) supplemented to contain 10 % FBS, 1% Penicillin-Streptomycin (Thermo Fisher Scientific #15140122) and 1% L-Glutamine (Thermo Fisher Scientific #A2916801). All cell lines were cultured at 37 °C and 5 % CO_2_. All cell lines were confirmed to be negative for mycoplasma contamination.

### Viruses

#### Influenza A/PR/8/34 virus

Influenza A/PR/8/34 virus (IAV PR8) stocks were generated using 9-day-old embryonated chicken eggs (Charles River #10100335). Eggs were incubated over night at 37 °C and candled by holding the eggs directly against a light source to identify an inoculation site without any veins above the air sac. The egg was inoculated with 1,000 plaque forming units (PFU) diluted in Dulbecco’s phosphate buffered saline (PBS, Thermo Fisher Scientific #14190144) + 1% BSA (Sigma Aldrich #A9576) using a 18G needle. Eggs were incubated for 2 days at 37 °C and transferred to 4 °C for 2 hours. Next, eggs were opened with a sterile spoon and a scoopula was used to push down the embryo and the allantoic fluid was collected. The allantoic fluid was centrifuged using an Allegra X-12R (Beckman Coulter) at 205 G for 5 min, the supernatant was transferred into a clean falcon tube and stored at -80 °C.

#### VSV-GFP

GFP-tagged vesicular stomatitis virus (VSV-GFP) ^53^ was generated by infecting 80% confluent BHK-21 cells (ATCC #CCL-10) at a multiplicity of infection (MOI) of 0.01 PFU/cell. Cells were cultured at 37 °C and cell culture supernatant was harvested after 2 days, clarified by centrifugation at 1,850 G for 5 min, and filtered through a 0.22 µm membrane and stored at -80 °C.

#### SARS-CoV-2 strain MA10

Severe acute respiratory syndrome coronavirus 2 (SARS-CoV-2) strain MA10 ^20^ was generously provided by Ralph Baric (University of North Carolina at Chapel Hill). A P1 stock was amplified in VeroE6 cells obtained from the ATCC that were engineered to stably express TMPRSS2 (VeroE6_TMPRSS2_). To generate a P2 working stock, VeroE6_TMPRSS2_ cells were infected at a multiplicity of infection (MOI) of 0.1 PFU/cell and incubated at 37 °C for 4 days. The virus-containing supernatant was harvested, clarified by centrifugation at 1,850 G for 5 min for 10 min, and filtered through a 0.22 μm membrane and stored at -80 °C.

### Plaque assay

#### Influenza A/PR/8/34 virus

IAV PR8 was titrated by plaque assay on MDCK cells (ATCC #CCL-34) in 6-well format. Briefly, 5×10^5^ cells per well were plated the day prior, medium was aspirated, and cells were washed with PBS (Thermo Fisher Scientific #14190144) before 500 µL of a serial dilution of viral inoculum in infection medium (RPMI, 0.1 % fetal bovine serum (FBS), 1 % Penicillin-Streptomycin, 0.3 % BSA, 1 µg/mL tosylamido phenylalanyl chloromethyl ketone (TPCK)-treated trypsin) was added. Cells were incubated for 60 min at 37 °C and plates were moved every 20 min to prevent drying out of the wells. The initial inoculum was removed, 2 mL of overlay were added per well (25 mL 2x DMEM, 0.05 mL FBS, 15 mL 2% oxoid agar (Thermo Fisher Scientific #LP0011B), 0.5 mL 30 % BSA, 0.5 mL 1 % diethylaminoethyl (DEAE)-dextran, 0.75 mL sodium bicarbonate (NaHCO_3_), 0.05 mL 1mg/mL TPCK trypsin, 8.15 mL water) and cells were incubated at 37 °C for 2 days. Cells were fixed by adding 2 mL 4 % paraformaldehyde (Sigma Aldrich #F8775) for 1 hour, the overlay was removed, and cells were permeabilized by incubating with 1 mL 0.1% Triton X-100 (Sigma Aldrich #93443) in PBS for 10 min at room temperature. Cells were washed 2 times with PBS, and incubated with 500 µL of anti-IAV nucleoprotein (NP) antibody (1:3000, Sigma Aldrich #MAB8257) in 5 % goat serum (Jackson Immuno Research #005-000-121) for 1 hour at 37 °C. Cells were washed 2 times with 0.05 % Tween-20 (Sigma Aldrich P1379) in PBS, and incubated with 500 µL anti-mouse-horseradish peroxidase (HRP) antibody (1:1000, Jackson Immuno Research #115-035-146) in 0.05 % Tween-20 in PBS at 37 °C for 1 hour. Next, cells were washed 2 times with 0.05 % Tween-20 in PBS, add 500 µL KPL TrueBlue Peroxidase Substrate (Seracare #5510-0052) and incubated for 1 min at room temperature. Cells were washed with PBS and viral titer was quantified by enumerating foci.

#### VSV-GFP

VSV-GFP was titrated by plaque assay on BHK-21 cells (ATCC #CCL-10) in 6-well format. Briefly, 5×10^5^ cells per well were plated the day prior, medium was aspirated and 500 µL before a 10-fold serial dilution of viral inoculum was added. Cells were incubated for 60 min at 37 °C and plates were moved every 20 min to prevent drying out of the wells. The initial inoculum was removed, 2 mL of overlay were added per well (25 mL 2x DMEM, 10 mL FBS, 15 mL 2% oxoid agar) and incubated for 2 days at 37 °C. Cells were fixed by adding 2 mL 4 % paraformaldehyde (Sigma Aldrich #F8775) for 1 hour at room temperature. The overlay was removed, and virus was quantified by enumerating GFP-positive foci and/or upon adding crystal violet solution (1.25 % crystal violet, 20 % methanol in distilled water) for 15 min at room temperature.

#### SARS-CoV-2 strain MA10

SARS-CoV-2 strain MA10 was titrated by plaque assay on VeroE6 cells obtained from Ralph Baric (University of North Carolina at Chapel Hill) that stably express TMPRSS2 (VeroE6_UNC/TMPRSS2_) (referred to as VeroE6_UNC_) in 6-well format. Briefly, 4×10^5^ cells were plated the day prior, medium was aspirated and 500 µL of a serial 10-fold virus dilutions in Opti-MEM were added. Cells were incubated for 90 min 37 °C and plates were moved every 20 min to prevent drying out of the wells. The initial inoculum was removed, 2 mL overlay were added per well (DMEM containing 10 % FBS with 1.2 % microcrystalline cellulose (Avicel)). Cells were incubated for four days at 33 °C, followed by fixation with 7 % formaldehyde and crystal violet staining for 1 hour. The overlay was removed, and virus was quantified by enumerating plaques. All SARS-CoV-2 MA10 experiments were performed in a biosafety level 3 laboratory. To verify SARS-CoV-2 MA10 identity and test for unwanted mutations, RNA from virus stocks was purified using TRIzol Reagent (Thermo Fisher Scientific, #15596026). Briefly, 200 µL of each virus stock was added to 800 µL TRIzol Reagent, followed by 200 µL chloroform, which was then centrifuged at 12,000 G for 5 min. The upper aqueous phase was moved to a new tube, mixed with an equal volume of isopropanol, and then added to an RNeasy Mini Kit column (QIAGEN, #74014) to be further purified following the manufacturer’s instructions. Viral stocks were subsequently confirmed via next generation sequencing using libraries for Illumina MiSeq.

### Intranasal treatments and infections

For anesthesia, mice were intraperitoneally injected with a mixture of ketamine (80mg/kg; Zoetis, #54771-2013-1) and xylazine (8.8 mg/kg; Akorn, #07-808-1947) in PBS. After mice were sufficiently anesthetized, 30µL of inoculum was applied to one nostril. Mice were monitored until they regained consciousness.

For infections, mice were inoculated with 6,000 PFU of SARS-CoV-2 MA10, 200 or 50 PFU of influenza A/PR/8/34 virus for LD50 and sublethal infections, respectively.

### Isolation of bronchoalveolar lavage fluid

Mice were euthanized and the trachea was carefully exposed. A 18G catheter equipped with a 3-way cock stop valve was inserted into the trachea. Bronchoalveolar lavage fluid (BALF) was obtained by flushing the airways with 5x 1mL of sterile PBS containing 2mM ethylenediaminetetraacetic acid (EDTA). BALF was stored on ice at all times until further use.

### Alveolar macrophage culture

Isolated BALF was counted, spun at 500 G for 5 minutes at 4 °C and resuspended in medium to reach the desired cell concentration.

For short-term culture, cells were resuspended and cultured in RPMI medium (Thermo Fisher Scientific #11875093) supplemented to contain 10% fetal bovine serum (FBS), 1% Penicillin-Streptomycin (Thermo Fisher Scientific #15140122) and 1% L-Glutamine (Thermo Fisher Scientific #A2916801). Cells were incubated over night, washed with PBS the next day and fresh medium containing the respective stimuli was added.

Long-term culture of alveolar macrophages was carried out as previously described ^25^. Briefly, cells were resuspended and cultured in RPMI medium (Thermo Fisher Scientific #11875093) supplemented to contain 10 % fetal bovine serum (FBS), 1 % Penicillin-Streptomycin (Thermo Fisher Scientific #15140122), 1 % L-glutamine (Thermo Fisher Scientific #A2916801), 30 ng/mL granulocyte-macrophage colony-stimulating factor (GM-CSF) (Peprotech #315-03), 10 ng/mL transforming growth factor (TGF)-β1 (Peprotech #100-21), 1 µM Rosiglitazone (Sigma Aldrich #R2408) and 50 µg/mL Gentamicin (Thermo Fisher Scientific #15750060). Cells were cultured in standard tissue culture vessels, medium was changed every 3 days and split when they reached 70-80 % confluency. Typical alveolar macrophage cultures consist of adherent and suspended cells. For splitting, culture supernatants were collected, cells were rinsed with Dulbecco’s phosphate buffered saline (PBS, Thermo Fisher Scientific #14190144). To detach adherent cells, ESGRO Complete Accutase (Sigma Aldrich #SF006) was added and cells were incubated at 37 °C for 5 to 10 minutes until cells were detached. Cells were completely detached and resuspended in medium by pipetting. Cells were counted on a Countess automated cell counter (Invitrogen #c10281), spun at 500 G for 5 minutes at 4 °C and resuspended in medium to reach the desired cell concentration and plated into a new cell culture vessel.

### *In vitro* polyIC treatment of alveolar macrophages

For all experiments, endotoxin-free polyinosinic-polycytidylic acid (polyIC, InvivoGen #tlrl-pic) was used. Stimulation of alveolar macrophages with polyIC was done at 10 µg/mL for 24 hours (establishment of innate immune memory) or 6 hours (acute response or recall response).

To establish innate immune memory in alveolar macrophages, cells were treated with 10 µg/mL polyIC for 24 hours. Cells were washed with PBS (Thermo Fisher Scientific #14190144) and fresh medium was added. 2 days after washing, fresh medium was added and/or cells were split if necessary. 4 days after washing, cells were seeded into 24-well or 96-well plates. Cells were re-stimulated the next day with polyIC for 6 hours (RT-qPCR, RNA-seq or immunofluorescence) or 24h (immunofluorescence).

### VSV-GFP infection of alveolar macrophages

VSV-GFP infection of alveolar macrophages was carried out at a MOI of 0.01 PFU/cell. Medium was aspirated from cell cultures and medium containing VSV-GFP at the desired dilution and 1 µg/mL Hoechst (Thermo Fisher Scientific #33342) was added. After 24 hours, medium was aspirated and cells were fixed for 10 min in 4 % paraformaldehyde (Sigma Aldrich #F8775) at room temperature, washed with 200 µL PBS (Thermo Fisher Scientific #14190144) and stored in 200 µL at 4 °C until imaging.

### Alveolar macrophage transfer *in vivo*

Anesthetized mice were intranasally (i.n.) treated with 2×25 µL clodronate liposomes (Liposoma #C-005) 3 days prior to alveolar macrophage transfer. Alveolar macrophages were isolated and expanded *in vitro* as described above. Cells were treated with 10 µg/mL polyIC (InvivoGen #tlrl-pic) or PBS (Thermo Fisher Scientific #14190144) and 1 day prior to transfer. At the day of transfer, cells were detached and washed 2 times with 50 mL PBS and counted and concentration was adjusted. Each mouse was i.n. transferred 700,000 cells in 30 µL PBS. Mice were allowed to recover for 5 days and were subsequently challenged with 200 PFU (LD50) of influenza A virus strain PR8 and body weight loss was monitored.

### RNA isolation

Cells or tissue samples were collected in 1 mL TRIzol Reagent (Thermo Fisher Scientific #15596026). Tissues were homogenized using glass beads (BioSpec Products #11079110) and a MagNA Lyser (Roche Diagnostics) at 6,000 rpm and 30 seconds. Total RNA was isolated according to the TRIzol/chloroform extraction protocol of the Ambion PureLink RNA Mini Kit (Thermo Fisher Scientific #12183025). RNA concentration was determined on a NanoDrop Instrument (Thermo Fisher Scientific) and RNA was stored at -80 °C.

### cDNA synthesis

cDNA was synthetized using the RevertAid First Strand cDNA Synthesis Kit and random hexamer primers (Thermo Fisher Scientific #K1622) according to the manufacturer’s instructions.

### qPCR

Quantitative PCRs (qPCR) were run and analyzed on a QuantStudio3 Real-Time PCR System (Thermo Fisher Scientific) according to the PowerUP SYBR Green (Thermo Fisher Scientific #25741) protocol with 1.5 to 15 ng cDNA input. Expression levels of *Eef1a* (5’-GCAAAAACGACCCACCAATG-3’, 5’-GGCCTGGATGGTTCAGGATA-3’), *Ifit1* (5’TTACAGCAACCATGGGAGAGAATG-3’, 5’-GGAACTGGACCTGCTCTGAGATTC-3’), *Ifitm3* (5’-GCCTACTCCGTGAAGTCTAGGG-3’, 5’-CCAAGGTGCTGATGTTCAGGC-3’), *Bst2* (5’-TGTAGAGACGGGTTGCGAGC-3’, 5’-CTCCTGAAGGGTCACCACGG-3’), *SARS-CoV-2 N* (5’-TAATCAGACAAGGAACTGATTA-3’, 5’-CGAAGGTGTGACTTCCATG-3’) and *PR8 M* (5’-CATGGAATGGCTAAAGACAAGACC-3’, 5’-CCATTAAGGGCATTTTGGACA-3’) were determined using a standard qPCR protocol (step 1: 20 sec at 95 °C, step 2: 1 sec at 95 °C, step 3: 20 sec at 60 °C, step 4: go to step 2 and repeat 40x, step 5: 1 sec at 95 °C, step 6: 20 sec at 60 °C, step 7: ramp down up to 95 °C at +0.15 °C/s, step 8: 1 sec 95 °C, step 9: end).

### Flow cytometry

Bronchoalveolar lavage fluid (BALF) was centrifuged at 500 G for 5 minutes at 4 °C. For bone marrow (BM), red blood cells (RBC) were lysed by incubating cells with RBC Lysis Buffer (Biolegend #420301) for 3 min at room temperature. To stop RBC lysis, 1 mL PBS (Thermo Fisher Scientific #14190144) was added, and cells were centrifuged at 500 G for 5 minutes at 4 °C. Supernatant was aspirated down to 100 µL, cells were resuspended and transferred to a V-bottom 96-well plate. Plates were centrifuged at 500 G for 5 minutes at 4 °C. Supernatant was aspirated and cells were resuspended in 25 µL FcR block per well (200x, Biolegend #101320) diluted in FACS buffer (2 % FBS, 2 mM EDTA, 10g/L NaN_3_) and incubated at room temperature for 15 min. This step was omitted for BM samples. Next, 25 µL 2x surface staining mix diluted in FACS buffer were added per well and cells were incubated for 30 min at 4 °C. For BM samples, 1x surface staining mix diluted in FACS buffer was added. Cells were washed with 150 µL PBS per well and resuspended in 100 µL viability dye. Cells were incubated for 15 min at room temperature and washed twice with 100 µL FACS buffer per well before fixation with 4 % paraformaldehyde (Sigma Aldrich #F8775) for 10 min at room temperature. Samples were washed once with 200 µL FACS buffer and resuspended in 150-250 µL FACS buffer and stored at 4 °C until acquisition.

For BALF surface staining, anti-F4/80 (BV421, clone BM8, Biolegend #123137), anti-CD4 (BV510, clone GK1.5, Biolegend #100449), anti-CD11c (BV605, clone N418, Biolegend #117333), anti-CD11b (BV650, clone M1/70, Biolegend #101259), anti-CD69 (BV711, clone H1.2F3, Biolegend #104537), anti-CD44 (FITC, clone IM7, Biolegend #103005), anti-SiglecF (PerCP-Cy5.5, clone S17007L, Biolegend #155531), anti-CD3 (PE-Dazzle, clone 17A2, Biolegend #100245), anti-CD62L (PE-Cy5, clone MEL-14, Biolegend #104410), anti-CD103 (PE-Cy7, clone QA17A24, Biolegend #156905), anti-Ly6G (APC, clone 1A8, Biolegend #127613), anti-CD8a (AF647, clone 53-6.7, Biolegend #100724), anti-CD45 (AF700, clone I3/2.3, Biolegend #147715), anti-NK1.1 (APC-Cy7, clone S1701D, Biolegend #156509), anti-CD45.2 (APC, clone 104, Biolegend #109813), anti-SiglecF (APC-Cy7, clone S17007L, Biolegend #155531), anti-CD11c (Pacific Blue, clone N418, Biolegend #117321) were used.

For BM surface staining, Biotin anti-mouse Lineage Panel (Biolegend #133307), anti-CD150 (BV650, clone TC15-12F12.2, Biolegend #115932), anti-CD117 (BV785, clone 2B8, Biolegend #105841), anti-CD48 (PerCP-Cy5.5, clone HM48-1, Biolegend #103421), anti-CD135 (PE, clone A2F10, Biolegend #135305), anti-Sca1 (PE-Dazzle, clone E13-161.7, Biolegend #122527), anti-CD16/32 (PE-Cy5, clone S17011E, Biolegend #156617), anti-CD34 (PE-Cy7, clone SA376A4, Biolegend #152217), anti-CD127 (APC, clone S18006K, Biolegend #158205), anti-CD115 (APC-Fire750, clone AFS98, Biolegend #135535), FITC Streptavidin (Biolegend #405201), BV421 Streptavidin (Biolegend #405226) were used.

For dead cell exclusion, Zombie Green Fixable Viability Kit (Biolegend #423111), Zombie Violet Fixable Viability Kit (Biolegend #423114) and LIVE/DEAD Fixable Blue dead Cell Stain Kit (Thermo Fisher Scientific #L23105) were used.

### PKH26 labeling

Mice were anesthetized and intranasally treated with a 1:50 dilution of LumiTrace PKH26 (Lumiprobe #13201). 5 days after treatment, mice were intranasally challenged with virus.

### Histological analyses

After euthanasia, murine lung tissues were excised and placed in 10 mL 10% neutral buffered formalin (Fischer Scientific #SF100-4) for 48 hours. Samples were subsequently transferred to 70% ethanol for processing in a Leica ASP300 following a one hour, 13 step program. Samples were embedded in paraffin using standard orientation procedures. Five-micron tissue sections were collected onto Plus slides (Fisher Scientific #22-042-924), air-dried, and stored at room temperature prior to use. Each sample was histochemically stained with hematoxylin-eosin.

Chromogenic immunohistochemistry (CIHC) was performed using unconjugated polyclonal rabbit anti-SARS CoV-2 Nucleocapsid protein (GeneTex Cat# GTX135357, Lot# 43979, RRID: AB_2868464). Lungs from SARS-CoV-2 recovered and naïve animals were sectioned on to the same slide and used for antibody optimization. Initial optimization testing determined antigen retrieval requirements at fixed concentration. Subsequent optimization manipulated concentration and or incubation to establish final protocol parameters. Negative controls consisted of primary antibody substituted with antibody diluent. All immunohistochemistry was performed on a Ventana Medical Systems Discovery Ultra platform using Ventana’s reagents and detection kits unless otherwise noted. In brief, sections were deparaffinized online. Rabbit anti-SARS-CoV-2 Nucleocapsid protein was diluted 1:200 (1.6 ug/ml) and incubated for 3 hours at room temperature and with goat anti-rabbit horseradish peroxidase conjugated for 8 minutes followed by DAB detection. Slides were washed in distilled water, counterstained with hematoxylin, dehydrated through graded alcohols, cleared in xylene and mounted with synthetic permanent media. Appropriate positive and negative controls were included with the study sections.

### Immunofluorescence

Immunofluorescence experiments were performed in black wall 96-well plates (Corning #3904). At the respective time point, cells were fixed for 10 min in 4 % paraformaldehyde (Sigma Aldrich #F8775) at room temperature and washed 2 times with 200 µL PBS (Thermo Fisher Scientific #14190144). Next, cells were incubated with 100 µL ice cold methanol (Sigma Aldrich #34860) for 10 min at -20 °C. After washing with 200 µL PBS, 50 µL of PBS containing 0.5% Triton X-100 (Sigma Aldrich #93443) was added for 5 minutes and then blocked with 50 µL 5% goat serum (Jackson Immuno Research #005-000-121) in PBS for 30 min at room temperature. Primary antibodies were added in 35 µL per well and incubated for 1 hour at room temperature. Cells were washed 3 times with 200 µL PBS for 4 min before secondary antibodies and 1 µg/mL Hoechst (Thermo Fisher Scientific #33342) were added in 35 µL per well and incubated for 1 hour at room temperature. Afterwards, cells were washed 3 times with 200 µL PBS for 4 min and stored in 200 µL PBS at 4 °C until imaging. Cells were imaged on a BioTek Cytation 7 or a Perkin Elmer Operetta CLS instrument.

### Image analyses

Images were analyzed and fluorescent intensities quantified using CellProfiler version 4.2.1 ^54^. Subsequent analyses were performed using RStudio version 2023.03.0+386.

### Cytokine profiling

Cytokines were quantified using Luminex xMAP technology for multiplexed quantification of 45 Mouse cytokines, chemokines and growth factors offered by Eve Technologies Corp. (Calgary, Alberta, Canada). According to Eve Technologies, the multiplexing analysis was performed using the Luminex™ 200 system (Luminex, Austin, TX, USA). Forty-five markers were simultaneously measured in the samples using Eve Technologies’ Mouse Cytokine 44-Plex Discovery Assay which consists of two separate kits; one 32-plex and one 13-plex (Sigma, Burlington, Massachusetts, USA). The assay was run according to the manufacturer’s protocol. The 32-plex consisted of Eotaxin, G-CSF, GM-CSF, IFNγ, IL-1α, IL-1β, IL-2, IL-3, IL-4, IL-5, IL-6, IL-7, IL-9, IL-10, IL-12(p40), IL-12(p70), IL-13, IL-15, IL-17, IP-10, KC, LIF, LIX, MCP-1, M-CSF, MIG, MIP-1α, MIP-1β, MIP-2, RANTES, TNFα, and VEGF. The 13-plex consisted of 6Ckine/Exodus2, Erythropoietin, Fractalkine, IFNβ-1, IL-11, IL-16, IL-20, MCP-5, MDC, MIP-3α, MIP-3β, TARC, and TIMP-1. Assay sensitivities of these markers range from 0.3 to 30.6 pg/mL for the 45-plex. Individual analyte sensitivity values are available in the Sigma MILLIPLEX® MAP protocol.

### Bulk transcriptomics

RNA was isolated as described above and were submitted to MedGenome Inc. (Foster City, California, USA). Sequencing libraries were prepped using the Takara SMARTer Stranded Total RNA-Seq Kit v3 or the Illumina TrueSeq RNA Library Prep Kit v2 and 150 bp paired-end sequencing was performed on an Illumina NovaSeq instrument. Reads were aligned to the Mus musculus genome mm10 GCRm38 (Fig 2) or GCRm39 (Fig 3) using STAR v. 2.7.10b. Read counts were normalized, statistically analyzed and differentially expressed genes between conditions identified using the DESeq2 package version 1.34.0 ^55^ in RStudio version 2023.03/0+386.

### Hierarchical clustering and gene ontology enrichment analyses

For hierarchical clustering of bulk RNA transcriptomics, lowly expressed genes were excluded (average FPKM across all conditions < 1) and only genes with an absolute log2 fold change greater than 1 and an adjusted *p*-value lower than 0.05 were considered. Significantly regulated genes of single cell RNA sequencing were identified using Seurat v4 ^22^. Gene ontology (GO) enrichment analyses for biological processes (BP) of identified genes of interest of gene clusters was performed using the clusterProfiler package version 4.2.2 ^56^ and hierarchical clustering was performed in in RStudio version 2023.03/0+386.

### Single cell RNA sequencing

#### Library preparation

Bronchoalveolar lavage fluid was centrifuged at 500 G for 5 minutes at 4 °C and supernatant was aspirated. Cells were resuspended in 300 µL PBS (Thermo Fisher Scientific #14190144), counted and adjusted to equivalent concentrations. Cell suspensions were processed for scRNA-seq with the 10x Chromium NextGEM Single-cell 3’ v3.1 kit according to manufacturer’s instructions. Each sample (naïve/PR8 or SARS2/PR8) was loaded to an individual 10x Chromium controller Chip G lane at a concentration for targeted recovery of 5,000 cells per lane. Barcode cDNA amplification was performed with 12 cycles of PCR. Following Bioanalyzer QC, libraries were pooled and sequenced on the Illumina NextSeq 500 instrument in paired end configuration (Read 1: 28 nt, Read 2: 55 nt).

#### Data processing

Sequencing data was mapped and quantified to per cell gene expression counts using CellRanger count (v5.0.0, 10x Genomics) with a mouse reference transcriptome (mm10) appended with annotations for IAV PR8 and SARS-CoV-2 MA10 (joint reference prepared with CellRanger mkref). Gene x cell matrices were further processed and analyzed with Seurat (v4.0.5) ^57^ in the R statistical framework (v4.0.3). After exclusion of several mouse RNAs (*Gm42418*, *Gm26917*, *AY036118*) associated with artifactual signals ^58,59^, quality thresholds were set based on data exploration, and cells with fewer than 500 RNA UMI counts or greater than 10 % mitochondrial RNA UMI counts were excluded from further analysis. Putative heterotypic doublets were identified with scDblFinder (v1.4.0) ^60^ and excluded. Putative erythrocytes, defined as cells with greater than 75% of RNA UMI counts composed of hemoglobin transcripts, were excluded.

#### Data analysis

SCtransform ^61^ (default parameters, with fraction of mitochondrial genes and Seurat cell cycle score difference as regression factors) was used for normalization and variable feature selection. Sample integration was performed with standard Seurat workflow; IAV PR8 genes (no SARS-CoV-2 reads were detected in any sample) and variable immune receptor genes (i.e. T and B cell receptor V, D, J genes) were excluded from integration and principal component dimensionality reduction. For dimensionality reduction, the first 60 principal components of the integrated dataset were used for UMAP generation, NearestNeighbor processing, and unsupervised graph-based clustering.

“Major cell groups” were annotated with SingleR ^62^ (v1.4.1, cluster mode and single cell mode for proliferating cells) and the Immunological Genome Project ^63^ reference datasets. Each major cell group was extracted, re-clustered, and further annotated (“Subpopulation level”, as in Figure 4) based on canonical marker gene expression patterns.

ISG gene set ^24^ expression scores were calculated per cell from log1p-normalized RNA counts using AddModuleScore() from Seurat v4.

### 10X Multiome sequencing

Bronchoalveolar lavage fluid from 3 animals per condition was pooled, centrifuged at 500 G for 5 minutes at 4 °C. The supernatant was aspirated, and dead cells were removed using the Dead Cell Removal Kit (Miltenyi #130-090-101) and LS columns (Miltenyi #130-122-729) according to manufacturer’s instructions. Afterwards, nuclei were isolated according to the 10x Genomics standard protocol (#CG000365 Rev C). Briefly, cells were centrifuged at 500 G for 5 minutes at 4 °C, supernatant was aspirated and 200 µL of lysis buffer (10 mM Tris-HCl, 10 mM NaCl, 3 mM MgCl_2_, 0.1 % Tween-20, 0.1 % IGEPAL CA630, 0.01 % digitonin, 1 % BSA, 1 mM DTT, 1 U/µL Sigma Protector RNase inhibitor in nuclease free water) was added and mixed by pipetting. Cells were incubated on ice for 3 min and 2 mL wash buffer (10 mM Tris-HCl, 10 mM NaCl, 3 mM MgCl_2_, 0.1 % Tween-20, 1 % BSA, 1 mM DTT, 1 U/µL Sigma Protector RNase inhibitor in nuclease free water) was added and mixed by inverting the tube. Samples were centrifuged at 500 G for 5 minutes at 4 °C, supernatant aspirated and nuclei washed two more times with 1 mL wash buffer. During the last wash step, nuclei were filtered through a 40 µm FLowMi filter (Sigma Aldrich #BAH136800040) into a new tube. After the last wash step, nuclei were resuspended in nuclei buffer (10x Genomics #PN-20000153), counted using a Countess 3 cell counter (Thermo Fisher Scientific #A49865) and concentration was adjusted to 3,000-5,000 nuclei/µL.

Nuclei were then processed using the Chromium Next GEM Single Cell Multiome ATAC + Gene Expression Reagent Bundle (10x Genomics #1000285) and Chromium Controller & Next GEM Accessory Kit (10x Genomics #1000202) following the manufacturer’s user guide (10x Genomics CG000338-Rev F). The single cell RNA and ATAC sequencing libraries were prepared using Dual Index Kit TT Set A (10x Genomics #1000215) and Single Index Kit N Set A (10x Genomics #1000212) respectively and sequenced on Illumina NovaSeq6000 platform.

#### Data Preprocessing

The Multiome data underwent preprocessing using the Cell Ranger ARC 1.0.0 pipeline and were aligned to the mm10 genome. Subsequently, the Cell Ranger output was processed using the Seurat Weighted Nearest Neighbor Pipeline. To eliminate low-quality cells, a filtering process was employed as previously described ^31^. Scrublet ^64^ was used on the RNAseq data to remove duplets. From the pooled data, low-quality cells from individual samples were filtered out. For the RNA-seq object, sctransform normalization was applied, followed by principal component analysis (PCA). The top 30 principal components were used for UMAP embedding and clustering. This process was repeated using 20 principal components.

Next, scATAC profiles from all samples were combined, and initial cell-type annotation was performed based on scRNA-seq annotations. Peaks were called for each cell type using MACS2 (version 2.1.2). Redundant peaks were removed based on the q-value obtained from MACS2. Using the resulting list of peak regions, the number of reads overlapping each peak window was determined for each unique cell barcode tag. This generated a matrix of peak-by-cell counts corresponding to ATAC reads within peaks for each cell profiled. High-quality cells were retained based on having a fraction of reads in peaks (FRiP) greater than 0.4 and a sequencing depth of more than 1000. The cells filtered out during this step were also removed from the scRNA object to ensure consistency across both modalities.

After quality control, the scRNA-seq object was reprocessed using sctranform, PCA, clustering, and UMAP embedding. Clusters were obtained from the scRNA-seq data using the default parameters of Seurat (30 PCs for PCA). The annotations for these clusters were finalized based on the expression of marker genes specific to distinct immune cell types. The same set of cells was retained in the scATAC-seq component of the Multiome data, and the annotations were transferred accordingly. The scATAC object was processed using the Signac pipeline, which involved TF-IDF normalization, singular value decomposition (SVD), UMAP embedding, and clustering.

#### Motif Analysis

To analyze motifs, position weight matrices (PWMs) from the JASPAR2020 database ^65^ and a motif occurrence matrix using the mm10 genome were added to the separate assay. Per-cell TF motif activity was calculated by employing the RunChromVAR function of Signac ^66^.

#### Differential gene expression analysis

For differential analysis of gene expression and TF activity, the FindMarker function of Seurat was used. The Wilcoxon test was employed for statistical testing.

### Human data generation and analyses

#### Sample acquisition

Study participants were enrolled at Weill Cornell Presbyterian Hospital during the initial infection wave of SARS-CoV-2 in New York City (spring to winter 2020) and were most likely infected by the original/non-variant strain of SARS-CoV-2. None of the patients had received a COVID-19 vaccine at time of blood collection. Peripheral blood mononuclear cells (PBMCs) and plasma was isolated and single-cell Multiome datasets were generated as previously described ^31^.

The classification of subjects into groups was based on the COVID-19 World Health Organization (WHO) Severity Classification. The groups included in this study were as follows: 1) healthy volunteer donors, and 2) recovered mild COVID-19 patients (with a WHO severity score of 1-2). To meet the inclusion criteria for each group, the following criteria were applied: 1) For healthy volunteer donors, individuals had to be free of any clinical symptoms related to COVID-19 at the time of blood collection. Negative SARS-CoV-2 PCR test results and/or seronegative status were also considered when available. 2) For recovered mild COVID-19 patients, individuals had to have a confirmed PCR test indicating SARS-CoV-2 infection, along with clinical symptoms of COVID-19 that did not require hospitalization. The prior infection status of both healthy volunteer donors and recovered mild COVID-19 patients was confirmed through SARS-CoV-2 serological testing after blood donation. Blood samples were collected using EDTA or sodium heparin-coated vacutainers and were kept on gentle agitation until processing. All blood samples were processed on the same day as collection. Information regarding age, sex, and comorbidities was obtained either through EPIC EHR records or, if not available, through a standardized form filled out at the time of donation.

---

**Figure S1:**
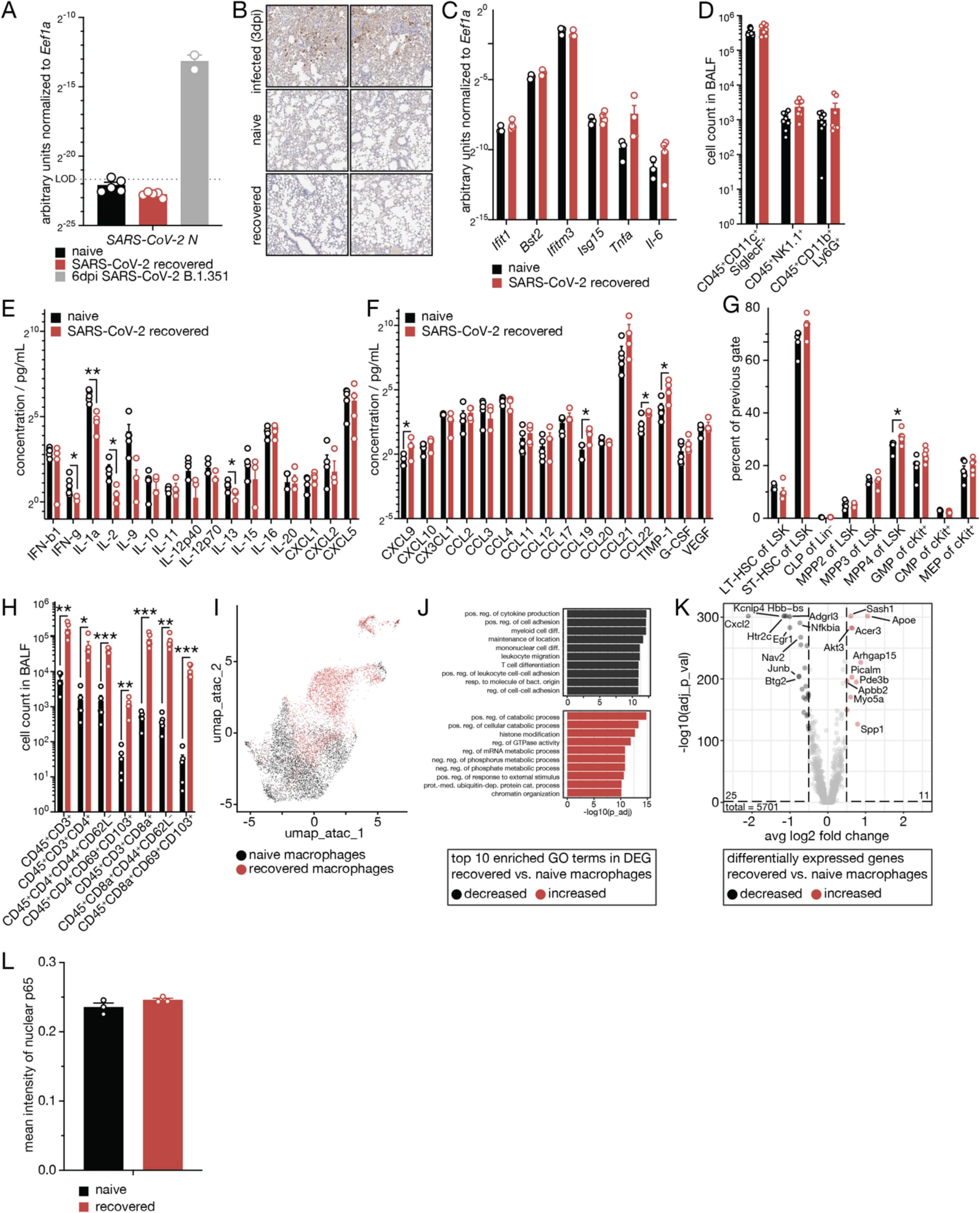
Characterization of naive and SARS-CoV-2-recovered animals. (A) RT-qPCR analyses of SARS-CoV-2 N transcript levels in lung tissue of naive, SARS-CoV-2-recovered or acutely infected (6 days post infection with SARS-CoV-2 strain B.1.351) C57BI/6J mice, n = 2-5. (B) Histologi­cal analyses of naive, SARS-CoV-2-recovered and acutely infected (3 days post infection with SARS-CoV-2 strain MA10). n = 5. (C) RT-qPCR analyses of Ifitl, Bst2, Ifitm3, Isg15, Tnfa and II-6 transcript levels in lung tissue of naive and SARS-CoV-2-recovered animals, n = 3-4. (D) Flow cytometric analyses of alveolar macro­phages (CD45+CD11 c+SiglecF+), NK cells (CD45+NK1.1 +) and neutrophils (CD45+CD11 b+Ly6G+) in broncho­alveolar lavage fluid (BALF) of naive and SARS-CoV-2-recovered animals, n = 7-8. (E-F) Cytokine profiling of BALF isolated from recovered and naive animals, n = 5. (G) Flow cytometric analyses of abundance of hemato­poietic progenitor cells (long-term hematopoietic stem cells (LT-HSC), short-term hematopoietic stem cells (ST-HSC), common lymphoid progenitor cells (CLP), multipotent hematopoietic progenitor cells (MPP), granulocyte-monocyte progenitor cells (GMP), common myeloid progenitor cells (CMP), megakaryocyte/eryth-roid progenitor cells (MEP)) in the bone marrow of naive and SARS-CoV-2-recovered animals, n = 5. (H) Flow cytometric analyses of T cell subsets in BALF of naive and SARS-CoV-2-recovered animals, n = 5. (I) UMAP clustering of the macrophage subset from single nuclei combined ATAC/RNA-sequencing data obtained from BALF of naive and SARS-CoV-2-recovered animals, n = 5. (J) Top 10 enriched gene ontology (GO) terms analyses of significantly regulated genes of recovered vs. naive macrophages (I). (K) Differentially expressed genes (DEG, absolute Iog2 fold change > 0.5) in macrophages (I) of recovered and naive animals. Top 10 significant DEG by fold change are labelled. (L) Quantification of mean fluorescent intensity of nuclear RELA (p65) in airway-resident macrophages isolated from naive and SARS-CoV-2-recovered animals, n = 3. Data are mean ± s.e.m. n values indicate the number of mice or replicates. For (A, C-H and L), statistics were calculated using Student’s t-test with Bonferroni correction when multiple comparisons were performed. For (K), statistical analysis was performed using Wilcoxon rank sum test with Bonferroni correction. For (J), hypergeometric p values were adjusted for multiple testing with Benjamini-Hochberg correction. *p < 0.05; **p < 0.01; ***p < 0.001.

**Figure S2:**
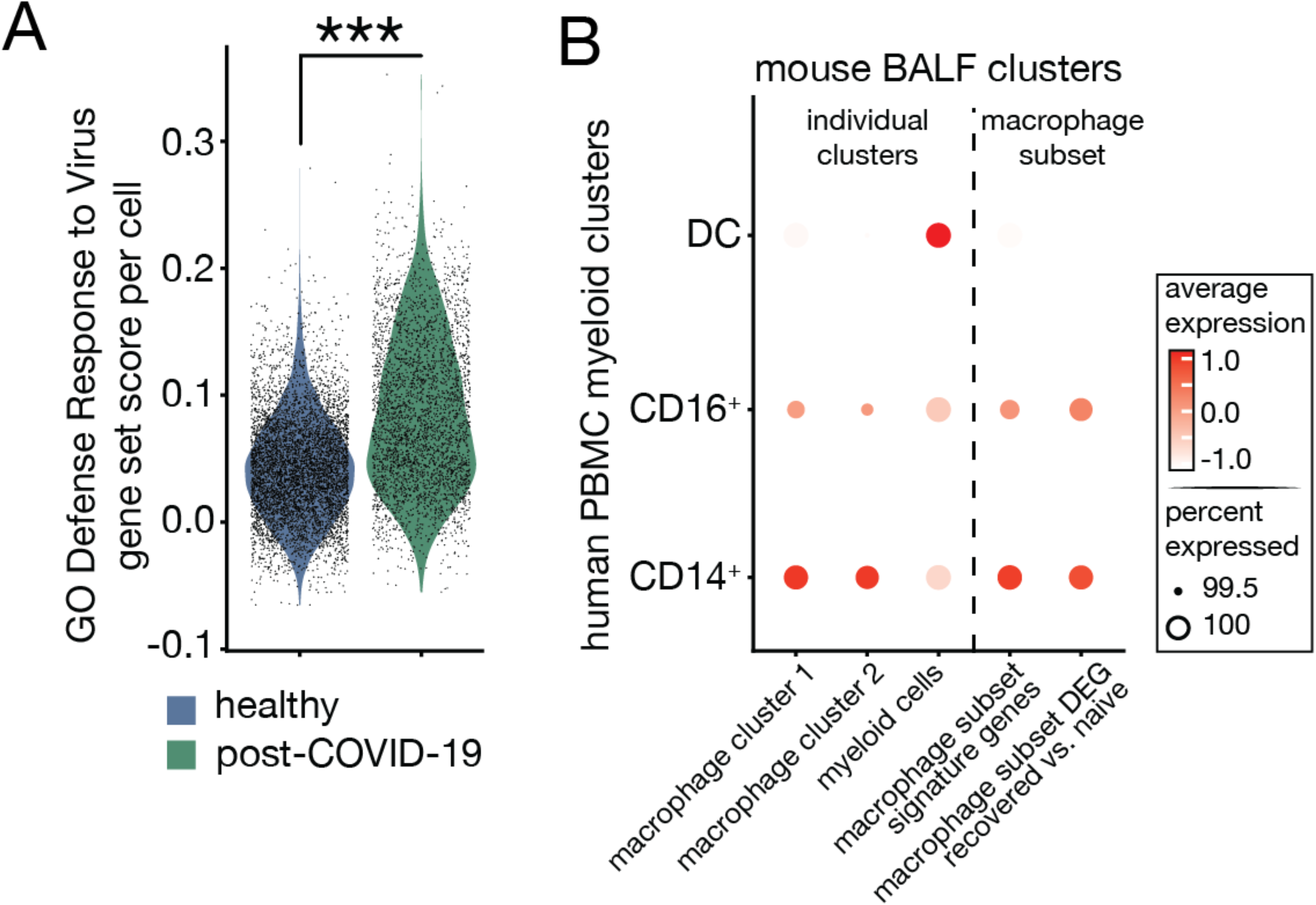
Characterization of naïve and SARS-CoV-2-recovered patient PBMCs. (A) Gene set expression score of GO: Defense Response to Virus across all recovered and healthy circulating human myeloid cells. (B) Expression of BALF cluster signature genes and macrophage subset (Figure 11) differentially expressed gene (DEG) modules in circulating human myeloid cells (0D14+ and 0D16+ monocytes and dendritic cells). Data are represented as violin plots or dotplot with each dot corresponding to one individual cell or cell cluster, respectively. For (A-B), statistical analysis was performed using Wilcoxon rank sum test with Bonferroni correction. *p < 0.05; **p < 0.01; ***p < 0.001.

**Figure S3:**
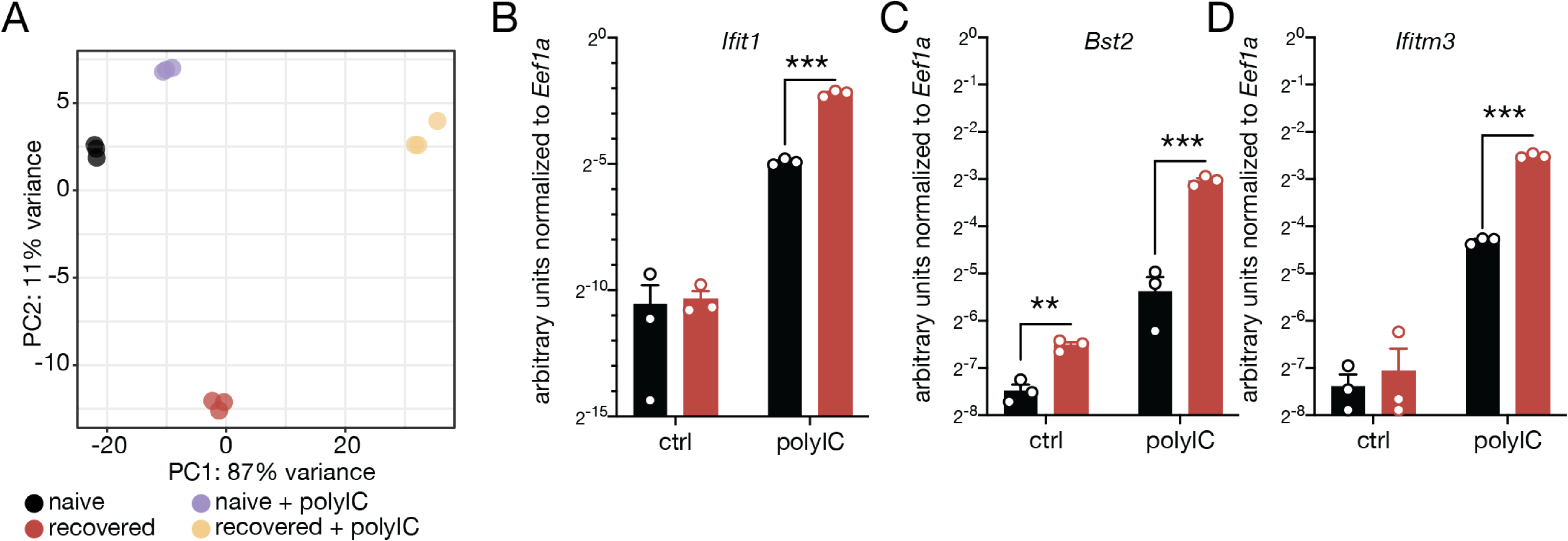
Characterization of secondary response of airway-resident macrophages isolated from naive and SARS-CoV-2-recovered animals. (A) Principal component analyses of transcriptomic profile of control- or polylC-stimulated airway-resident macrophages isolated from naive or SARS-CoV-2-recovered animals, n = 3. (B-D), RT-qPOR analyses of Ifitl (B), Ifitm3 (0) and Bst2 (D) transcript levels in control- or polylC-stimulated airway-resident macrophages isolated from naive or SARS-CoV-2-recovered animals, n = 3. Data are mean ±s.e.m. n values indicate the number of mice or replicates. For (B-D), statistics were calculated using Student’s t-test with Bonferroni correction when multiple comparisons were performed. *p < 0.05; **p < 0.01; ***p < 0.001.

**Figure S4:**
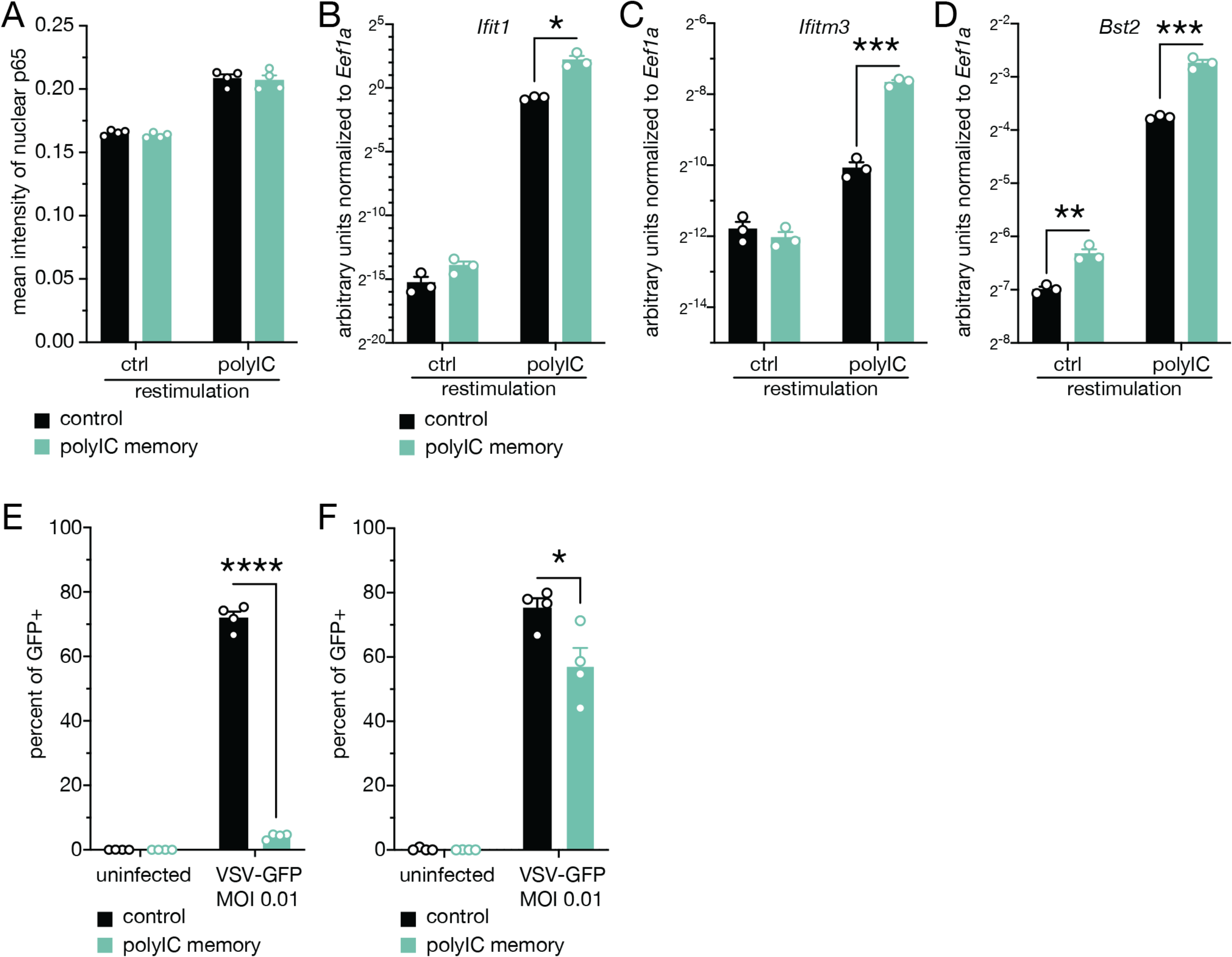
Characterization of secondary response of polylC-experienced alveolar macrophages in vitro. (A) Quantification of mean fluorescent intensity of RELA (p65) in control- or polylC-stimulated control- or polylC-experienced in vitro cultured alveolar macrophages after 6 hours, n = 4. (B-D), RT-qPCR of Ifitl (B), Ifitm3 (C) and Bst2 (D) transcript levels of polylC/polylC vs. control/polylC stimulated in vitro cultured alveolar macrophages 6 hours after re-stimulation, n = 3. (E-F), Percent of VSV-GFP-infected control- or polylC-stimulated control- or polylC-experienced in vitro cultured alveolar macro­phages after 5 (E) and 14 days (F). n = 4. Data are mean ± s.e.m. n values indicate the number of mice or replicates. For (A-F), statistics were calculated using Student’s t-test with Bonferroni correction when multiple comparisons were performed. *p < 0.05; **p < 0.01; ***p < 0.001.

**Figure S5:**
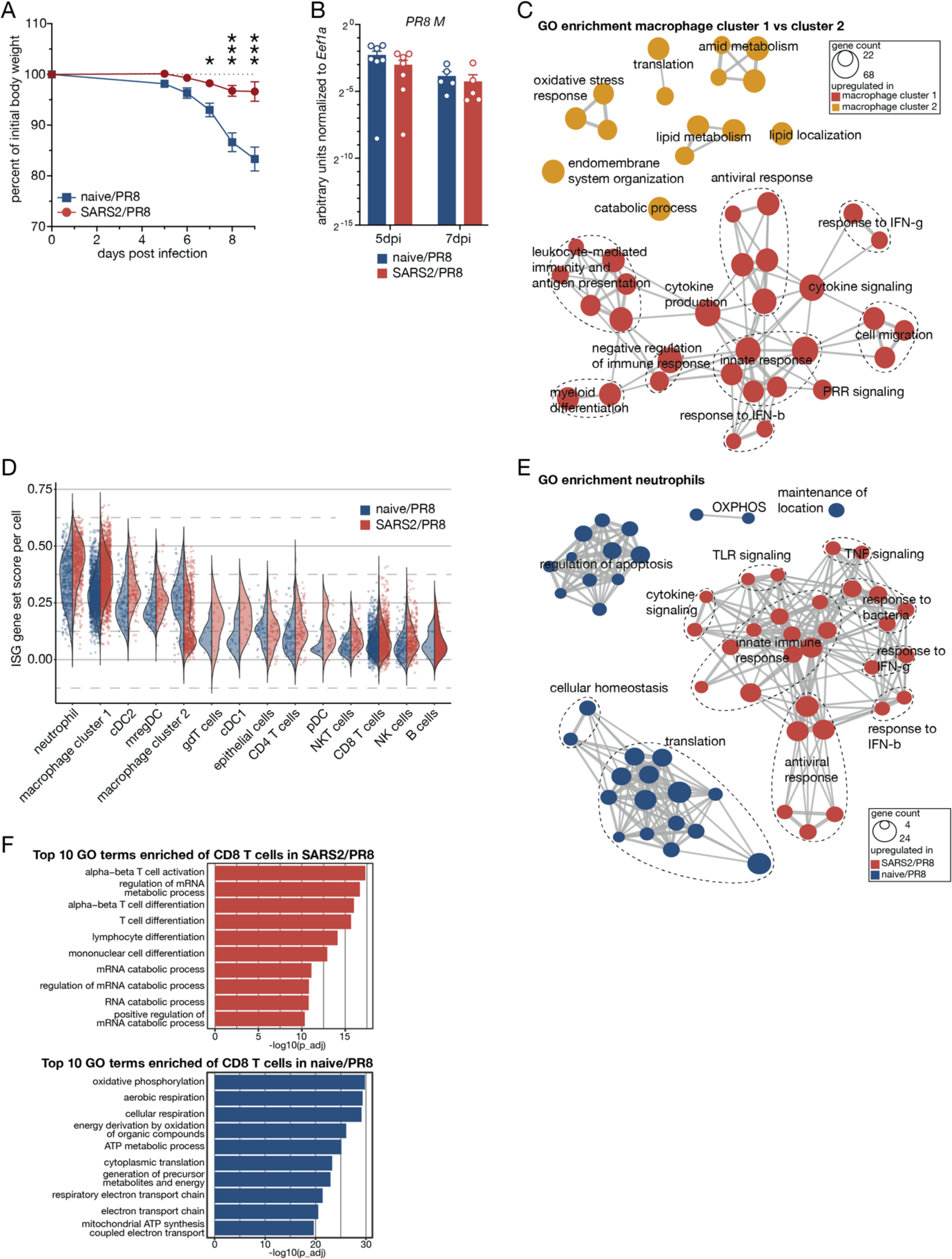
Characterization of pathology of naive and SARS-CoV-2-recovered animals superinfected with influenza A/PR/8/34 virus. (A) Body weights of naive and SARS-CoV-2-recovered animals infected with a sub-lethal dose of influenza A/PR/8/34 virus (naïve/PR8 or SARS2/PR8). n = 5-6. (B) RT-qPCR analyses of influenza A virus matrix protein (M1) transcript levels in naive/PR8 or SARS2/PR8 animals, n = 5-7. (C) Gene ontology (GO) enrichment analyses of differentially expressed genes (DEG) of macrophage cluster 1 vs. macrophage cluster 2 identified in single cell RNA-sequencing (scRNA-seq) of bronchoalveolar lavage fluid (BALF) in naive/PR8 or SARS2/PR8 animals at 7 days post PR8 infection. (D) Split violin plot of ISG gene set expression between any major cell populations and conditions identified by scRNA-seq (C). (E) GO enrichment analyses of DEG of neutrophils isolated from naïve/PR8 or SARS2/PR8 animals. Data are mean ± s.e.m. n values indicate the number of mice or replicates. (F) GO enrichment analyses of DEG of CD8 T cells isolated from naïve/PR8 or SARS2/PR8 animals. Data are mean ± s.e.m. n values indicate the number of mice or replicates. For (A), statistical analysis was performed using Two-Way ANOVA comparison with Bonferroni correction. For (B), statistics were calculated using Student’s t-test with Bonferroni correction when multiple comparisons were performed. For (C-F), DEG were identified using Wilcoxon rank sum test with Bonferroni correction. For (C-E and F), hypergeometric p values were adjusted for multiple testing with Benjamini-Hochberg correction. *p < 0.05; ∼p < 0.01; ***p < 0.001.

## References

1. Randolph, H. E. & Barreiro, L. B. Herd Immunity: Understanding COVID-19. Immunity 52, 737–741 (2020).

2. Naik, S. & Fuchs, E. Inflammatory memory and tissue adaptation in sickness and in health. Nature 607, 249–255 (2022).

3. Foster, S. L., Hargreaves, D. C. & Medzhitov, R. Gene-specific control of inflammation by TLR-induced chromatin modifications. Nature 447, 972–978 (2007).

4. Netea, M. G. et al. Defining trained immunity and its role in health and disease. Nat. Rev. Immunol. (2020) doi:10.1038/s41577-020-0285-6.

5. Naik, S. et al. Inflammatory memory sensitizes skin epithelial stem cells to tissue damage. Nature 550, 475–480 (2017).

6. Kaufmann, E. et al. BCG Educates Hematopoietic Stem Cells to Generate Protective Innate Immunity against Tuberculosis. Cell 172, 176–190.e19 (2018).

7. de Laval, B. et al. C/EBPβ-Dependent Epigenetic Memory Induces Trained Immunity in Hematopoietic Stem Cells. Cell Stem Cell 26, 657–674.e8 (2020).

8. Yao, Y. et al. Induction of Autonomous Memory Alveolar Macrophages Requires T Cell Help and Is Critical to Trained Immunity. Cell 175, 1634–1650.e17 (2018).

9. Kleinnijenhuis, J. et al. Long-lasting effects of BCG vaccination on both heterologous Th1/Th17 responses and innate trained immunity. J. Innate Immun. 6, 152–8 (2014).

10. Cirovic, B. et al. BCG Vaccination in Humans Elicits Trained Immunity via the Hematopoietic Progenitor Compartment. Cell Host Microbe 28, 322–334.e5 (2020).

11. Moorlag, S. J. C. F. M. et al. BCG Vaccination Induces Long-Term Functional Reprogramming of Human Neutrophils. Cell Rep. 33, 108387 (2020).

12. Garly, M. L. et al. BCG scar and positive tuberculin reaction associated with reduced child mortality in West Africa: A non-specific beneficial effect of BCG? Vaccine 21, 2782–2790 (2003).

13. Zhou, P. et al. A pneumonia outbreak associated with a new coronavirus of probable bat origin. Nature 579, 270–273 (2020).

14. Carabelli, A. M. et al. SARS-CoV-2 variant biology: immune escape, transmission and fitness. Nat. Rev. Microbiol. 21, 162–177 (2023).

15. Aegerter, H., Lambrecht, B. N. & Jakubzick, C. V. Biology of lung macrophages in health and disease. Immunity 55, 1564–1580 (2022).

16. Hashimoto, D. et al. Tissue-resident macrophages self-maintain locally throughout adult life with minimal contribution from circulating monocytes. Immunity 38, 792–804 (2013).

17. Zhu, B. et al. Uncoupling of macrophage inflammation from self-renewal modulates host recovery from respiratory viral infection. Immunity 54, 1200–1218.e9 (2021).

18. Zahalka, S. et al. Trained immunity of alveolar macrophages requires metabolic rewiring and type 1 interferon signaling. Mucosal Immunol. 15, 896–907 (2022).

19. Wang, T. et al. Influenza-trained mucosal-resident alveolar macrophages confer long-term antitumor immunity in the lungs. Nat. Immunol. 24, 423–438 (2023).

20. Leist, S. R. et al. A Mouse-Adapted SARS-CoV-2 Induces Acute Lung Injury and Mortality in Standard Laboratory Mice. Cell 183, 1070–1085.e12 (2020).

21. Dinnon, K. H. et al. SARS-CoV-2 infection produces chronic pulmonary epithelial and immune cell dysfunction with fibrosis in mice. Sci. Transl. Med. 14, (2022).

22. Hao, Y. et al. Integrated analysis of multimodal single-cell data. Cell 184, 3573–3587.e29 (2021).

23. Schep, A. N., Wu, B., Buenrostro, J. D. & Greenleaf, W. J. chromVAR: inferring transcription-factor-associated accessibility from single-cell epigenomic data. Nat. Methods 14, 975–978 (2017).

24. Schoggins, J. W. et al. A diverse range of gene products are effectors of the type i interferon antiviral response. Nature 472, 481–485 (2011).

25. Gorki, A.-D. et al. Murine Ex Vivo Cultured Alveolar Macrophages Provide a Novel Tool to Study Tissue-Resident Macrophage Behavior and Function. Am. J. Respir. Cell Mol. Biol. 66, 64–75 (2022).

26. Brandes, M., Klauschen, F., Kuchen, S. & Germain, R. N. A systems analysis identifies a feedforward inflammatory circuit leading to lethal influenza infection. Cell 154, 197–212 (2013).

27. Mitroulis, I. et al. Modulation of Myelopoiesis Progenitors Is an Integral Component of Trained Immunity. Cell 172, 147–161.e12 (2018).

28. Kaufmann, E. et al. BCG vaccination provides protection against IAV but not SARS-CoV-2. Cell Rep. 38, 110502 (2022).

29. Khan, N. et al. M. tuberculosis Reprograms Hematopoietic Stem Cells to Limit Myelopoiesis and Impair Trained Immunity. Cell 183, 752–770.e22 (2020).

30. Larsen, S. B. et al. Establishment, maintenance, and recall of inflammatory memory. Cell Stem Cell 1–17 (2021) doi:10.1016/j.stem.2021.07.001.

31. Cheong, J.-G. et al. Epigenetic Memory of COVID-19 in Innate Immune Cells and Their Progenitors. bioRxiv 2022.02.09.479588 (2022) doi:10.1101/2022.02.09.479588.

32. Murphy, J., Summer, R., Wilson, A. A., Kotton, D. N. & Fine, A. The prolonged life-span of alveolar macrophages. Am. J. Respir. Cell Mol. Biol. 38, 380–5 (2008).

33. Misharin, A. V et al. Monocyte-derived alveolar macrophages drive lung fibrosis and persist in the lung over the life span. J. Exp. Med. 214, 2387–2404 (2017).

34. Aegerter, H. et al. Influenza-induced monocyte-derived alveolar macrophages confer prolonged antibacterial protection. Nat. Immunol. 21, 145–157 (2020).

35. Li, F. et al. Monocyte-derived alveolar macrophages autonomously determine severe outcome of respiratory viral infection. Sci. Immunol. 7, (2022).

36. Wimmers, F. et al. The single-cell epigenomic and transcriptional landscape of immunity to influenza vaccination. Cell 1–21 (2021) doi:10.1016/j.cell.2021.05.039.

37. Rodríguez-Prados, J.-C. et al. Substrate fate in activated macrophages: a comparison between innate, classic, and alternative activation. J. Immunol. 185, 605–14 (2010).

38. Jha, A. K. et al. Network integration of parallel metabolic and transcriptional data reveals metabolic modules that regulate macrophage polarization. Immunity 42, 419–430 (2015).

39. Lercher, A., Baazim, H. & Bergthaler, A. Systemic Immunometabolism: Challenges and Opportunities. Immunity 53, 496–509 (2020).

40. Gu, H. et al. Vaccination induces rapid protection against bacterial pneumonia via training alveolar macrophage in mice. Elife 10, (2021).

41. Kamada, R. et al. Interferon stimulation creates chromatin marks and establishes transcriptional memory. Proc. Natl. Acad. Sci. U. S. A. 115, E9162–E9171 (2018).

42. Pulendran, B. & Ahmed, R. Translating innate immunity into immunological memory: implications for vaccine development. Cell 124, 849–63 (2006).

43. Ifrim, D. C. et al. Trained immunity or tolerance: Opposing functional programs induced in human monocytes after engagement of various pattern recognition receptors. Clin. Vaccine Immunol. 21, 534–545 (2014).

44. Novakovic, B. et al. β-Glucan Reverses the Epigenetic State of LPS-Induced Immunological Tolerance. Cell 167, 1354–1368.e14 (2016).

45. Roquilly, A. et al. Author Correction: Alveolar macrophages are epigenetically altered after inflammation, leading to long-term lung immunoparalysis. Nat. Immunol. 21, 962– 962 (2020).

46. Jurado-Camino, T. et al. Chronic lymphocytic leukemia: a paradigm of innate immune cross-tolerance. J. Immunol. 194, 719–27 (2015).

47. Jeljeli, M. et al. Trained immunity modulates inflammation-induced fibrosis. Nat. Commun. 10, 5670 (2019).

48. Ordovas-Montanes, J. et al. Allergic inflammatory memory in human respiratory epithelial progenitor cells. Nature 560, 649–654 (2018).

49. Lechner, A. et al. Macrophages acquire a TNF-dependent inflammatory memory in allergic asthma. J. Allergy Clin. Immunol. 149, 2078–2090 (2022).

50. Li, X. et al. Maladaptive innate immune training of myelopoiesis links inflammatory comorbidities. Cell 185, 1709–1727.e18 (2022).

51. Kimura, T. et al. Essential and non-redundant roles of p48 (ISGF3 gamma) and IRF-1 in both type I and type II interferon responses, as revealed by gene targeting studies. Genes Cells 1, 115–24 (1996).

52. Blight, K. J., McKeating, J. A. & Rice, C. M. Highly permissive cell lines for subgenomic and genomic hepatitis C virus RNA replication. J. Virol. 76, 13001–14 (2002).

53. Dalton, K. P. & Rose, J. K. Vesicular stomatitis virus glycoprotein containing the entire green fluorescent protein on its cytoplasmic domain is incorporated efficiently into virus particles. Virology 279, 414–21 (2001).

54. Stirling, D. R. et al. CellProfiler 4: improvements in speed, utility and usability. BMC Bioinformatics 22, 433 (2021).

55. Love, M. I., Huber, W. & Anders, S. Moderated estimation of fold change and dispersion for RNA-seq data with DESeq2. Genome Biol. 15, 550 (2014).

56. Wu, T. et al. clusterProfiler 4.0: A universal enrichment tool for interpreting omics data. Innov. 2, 100141 (2021).

57. Stuart, T. et al. Comprehensive Integration of Single-Cell Data. Cell 177, 1888–1902.e21 (2019).

58. Liu, Y. et al. Single-Cell Profiling Reveals Divergent, Globally Patterned Immune Responses in Murine Skin Inflammation. iScience 23, 101582 (2020).

59. Kimmel, J. C. et al. Murine single-cell RNA-seq reveals cell-identity- and tissue-specific trajectories of aging. Genome Res. 29, 2088–2103 (2019).

60. Germain, P.-L., Lun, A., Garcia Meixide, C., Macnair, W. & Robinson, M. D. Doublet identification in single-cell sequencing data using scDblFinder. F1000Research 10, 979 (2021).

61. Hafemeister, C. & Satija, R. Normalization and variance stabilization of single-cell RNA-seq data using regularized negative binomial regression. Genome Biol. 20, 296 (2019).

62. Aran, D. et al. Reference-based analysis of lung single-cell sequencing reveals a transitional profibrotic macrophage. Nat. Immunol. 20, 163–172 (2019).

63. Heng, T. S. P., Painter, M. W. & Immunological Genome Project Consortium. The Immunological Genome Project: networks of gene expression in immune cells. Nat. Immunol. 9, 1091–4 (2008).

64. Wolock, S. L., Lopez, R. & Klein, A. M. Scrublet: Computational Identification of Cell Doublets in Single-Cell Transcriptomic Data. Cell Syst. 8, 281–291.e9 (2019).

65. Fornes, O. et al. JASPAR 2020: update of the open-access database of transcription factor binding profiles. Nucleic Acids Res. 48, D87–D92 (2020).

66. Stuart, T., Srivastava, A., Madad, S., Lareau, C. A. & Satija, R. Single-cell chromatin state analysis with Signac. Nat. Methods 18, 1333–1341 (2021).

